# Deep learning from multiple experts improves identification of amyloid neuropathologies

**DOI:** 10.1101/2021.03.12.435050

**Authors:** Daniel R. Wong, Ziqi Tang, Nicholas C. Mew, Sakshi Das, Justin Athey, Kirsty E. McAleese, Julia K. Kofler, Margaret E. Flanagan, Ewa Borys, Charles L. White, Atul J. Butte, Brittany N. Dugger, Michael J. Keiser

## Abstract

Pathologists can have complementary assessments and focus areas when identifying and labeling neuropathologies. A standardized approach would ideally draw on the expertise of the entire cohort. We present a deep learning (DL) framework that consistently labels cored, diffuse, and cerebral amyloid angiopathy (CAA) neuropathologies using expert consensus. We collected 100,495 annotations, comprising 20,099 candidate neuropathologies from three institutions, independently annotated by five experts. We compared DL methods that learned the annotation behaviors of individual experts (AUPRC=0.67±0.06 cored; 0.48±0.06 CAA) versus those that reproduced expert consensus, yielding 8.9-13% improvements (AUPRC=0.73±0.03 cored; 0.54±0.06 CAA). Saliency mapping on neuropathologies illustrated how human expertise may progress from novice to expert. In blind prospective tests of 52,555 subsequently expert-annotated images, the models accurately labeled pathologies similar to their human counterparts (consensus model AUPRC=0.73 cored; 0.68 CAA).

## Introduction

Specific protein accumulations and neuroanatomic vulnerability define neurodegenerative diseases^1^. Studying these accumulations and accurately characterizing their presence, localization, and distribution improves understanding of pathophysiology^2^. Extracellular accumulations of amyloid beta (Aβ) into plaques are a pathological hallmark of Alzheimer’s disease. Aβ aggregates form diverse plaque morphologies, as well as deposits within blood vessels, termed cerebral amyloid angiopathy (CAA)^3,4^. These morphologies may change with disease severity and correlate to select clinical features^5,6^. Historically, however, the Consortium to Establish a Registry for Alzheimer’s Disease uses a semiquantitative measure of neuropathological phenotypes in its criteria^7^. Having more consistent, quantitative, and precise anatomic measures of Aβ aggregates would not only systematize and standardize research endeavors to better understand pathophysiology across institutions, but would likewise allow for detection of more subtle pathophysiological differences.

In many medical disciplines, there are inherent differences and areas of complementary expertise in evaluation. This becomes nuanced when differences in classification could possibly lead to differences in prognosis, treatment, and prevention measures^8–10^. Therefore, we posited that pathology assessment could benefit from more standardized approaches to improve quality and reliability, especially for imaging data^7,11^. In cases of diagnostic discrepancy, it can be difficult to determine what is objective truth. Certain medical professions, especially neuropathology, are also facing dwindling workforces despite high demand for their services^12^. Hence, some medical centers may not have pathologists to render interpretations or may lack bandwidth to collect multiple opinions on a case. Here, we present an automated framework of learning from multiple neuropathology experts to provide a more robust labeling co-pilot tool. We encapsulated this strategy into a DL model resilient to discrepancies among different expertises.

Additionally, pathological annotation can be a laborious and time-intensive process that reflects the unique training of the neuropathologist ^2,11,13^. In previous work, we automated a single expert’s annotations using DL^14^. This approach was validated by independent study with a different cohort^15^. However, individual bias of the expert remained, and it was not yet clear if these methods could scale across multiple experts, institutions, and data modalities—all of which are critical for assessing generalizability. In the current study, we augment five experts across a methodically and geographically diverse dataset.

Ground truth is difficult to ascertain for challenging pathology tasks. Guan and Hinton^16^ trained neural network ensembles to mimic individual doctors, and created a benchmark of ground-truth examples of retinal neuropathy diagnosis by adjudicating decisions among experts. Although an active discussion may be a judicial way to handle more difficult labeling, a consortium of experts is not always available, requires much time and labor, and presents confounding social factors that may bias labeling. Avoiding human adjudication and performing the task accurately and automatically at scale—while still utilizing diverse expert proficiency—may be optimal and time-efficient as a first-pass or a second-opinion assessment. Other approaches have assigned reliability scores to each expert, and used statistical procedures to estimate ground-truth^17–20^. We assessed complementary approaches, by both modeling individual learners and also augmenting a consensus-voting scheme among experts. The consensus models learned the final assessment using various voting thresholds, resulting in different levels of sensitivity and precision. We evaluated which of these two approaches was the most robust and performant—learning from individual assessments or learning from some consensus of experts. Finally, we built on the approach presented by Guan and Hinton^16^, and created ensembles that were robust to intentionally noisy information.

Although DL can potentially provide more standardized and quantitative assessments for pathology^21,22^, the complex mathematical functions learned can lack interpretability. There is a justifiable hesitancy to adopt such technology in high stakes scenarios without knowing how and why a model’s decisions are made^23,24^. Recent advances in interpreting complex DL models, primarily through the use of saliency mapping, begin to elucidate this decision-making process^25^. Employing such techniques, we found that DL models trained on individuals and on consensus annotations consistently learned relevant pathological features not explicitly given during training. The models’ feature detection was granular to the pixel level, and may help better understand features considered by human pathologists with different levels of expertise.

In addition to interpretability, we evaluated these models in a prospective research study. The DL models accurately adopted annotation patterns of their expert counterparts. We found that models trained from a consensus of experts were able to (1) reproduce consensus annotations, (2) prospectively filter and enrich for neuropathologies in new whole slide images (WSIs), (3) and exceed the performance of models trained from individual experts at the same tasks. We found that modeling a consensus of experts was performant even when evaluated on different benchmarks that were expected to favor individual models, advocating for the robustness of consensus learning. This methodology may be a powerful means to leverage diverse and complementary human expertise—among pathologists and more generally.

## Results

### We curated a multi-annotator dataset and found annotation differences among experts and consensus schemes

In prior work^14^, we developed a convolutional neural network (CNN) pipeline to automatically identify three different Aβ neuropathologies for a single expert. In this current study, we generalized this method to five experts (NP1-NP5) and two novice annotators (UG1 and UG2). Oftentimes, pragmatic DL applications are trained on data with limited diversity, resulting in inability to effectively generalize to data outside their training corpus^26^. To counter this, we validated our method using a dataset of 43 WSIs obtained from three different institutions, where each used different histological staining procedures (Methods). Supplementary Figure 1 contains demographic information on these 43 research patients.

We organized the study into two phases of data collection. In phase-one, we collected independent annotations from the seven annotators on the same 20,099 images derived from 29 WSIs. As in previous work, we color-normalized the WSIs^27^, identified candidate Aβ aggregates with conventional computer vision techniques, and center-cropped these candidates to provide 256 × 256 pixel images for final annotation (Methods). We arranged these candidates in a randomly shuffled but fixed ordering, and uploaded them to a custom web interface for expert annotation (Figure 1A). We recruited five neuropathology experts and two undergraduate novices from five different medical institutions to annotate the images. Annotators performed a multi-labeling task and independently labeled an image as any combination of “cored,” “diffuse,” and “CAA.” The annotators also had options of marking “negative,” “flag,” and “not sure,” but we did not use these to alter our image labels (Supplementary Figure 2). As in previous work, the diffuse class was most prevalent (Figure 1B).

**Figure 1.**
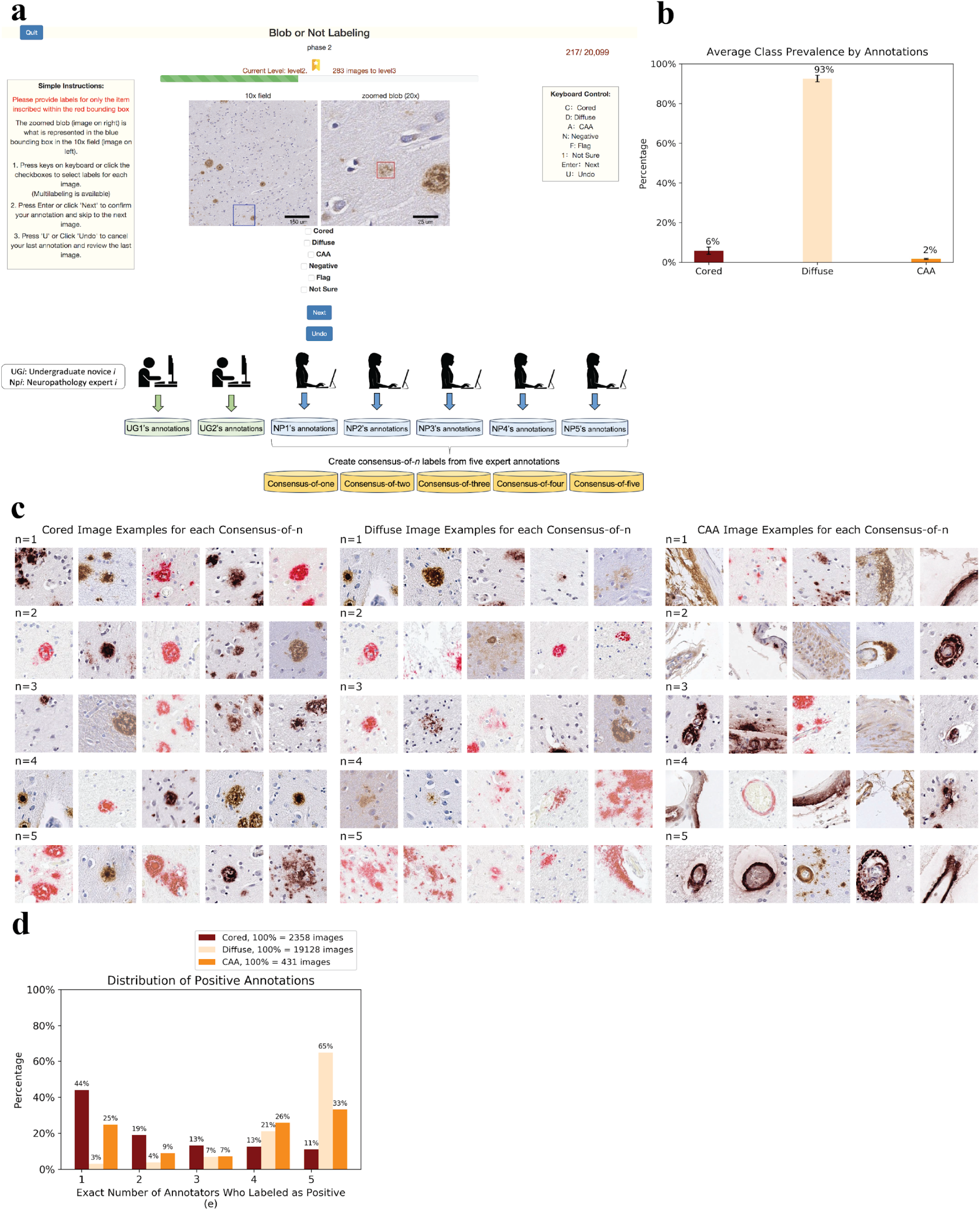
We curated annotations of Aβ neuropathologies from multiple experts, and found differing degrees of consensus. (a) Five experts (NP) and two undergraduate novices (UG) used a custom web portal for annotation. Each annotator labeled the same set of images in the same order. From the expert annotations, we constructed consensus-of-*n* labels (*n*=1 to *n*=5) for the same 20,099 images. (b) Average class distributions are consistent across the seven annotators. The y-axis plots average frequency, while the x-axis plots the Aβ class. (c) Representative images illustrating consensus-of-*n* strategies applied to each Aβ class, with rows progressing from top to bottom in order of increasing consensus. For a consensus-of-*n* image, at least *n* experts labeled the image as positive for the designated class. Each image was randomly and independently chosen from the set of images. (d) Positive annotation distributions differ by Aβ class. The x-axis plots the exact (not cumulative) number of annotators who gave a positive label. Hence, when *e*=1 and *e*=5 this is equivalent to a consensus-of-one and consensus-of-five respectively. For *e*=2, 3, or 4, this is *not* equivalent to an at-least-*n* consensus strategy. The y-axis plots the frequency. Each class has a different count of total positive labels (indicated in the legend). This total count represents the total number of images with at least one expert identifying the class. Each image may have multiple classes present.

To draw on the complementary proficiencies of multiple experts, we introduced the concept of a “consensus-of-*n*” strategy to label pathologies. In the consensus-of-*n* strategy, an image was labeled positive for plaque *p* if at least *n* experts positively labeled the image as presenting *p*. Otherwise, the image was assigned as negative for *p*. A consensus-of-one was the most permissive strategy, in which only one expert needed to positively identify *p* for this image to be labeled as positive for *p*. A consensus-of-one (logical set union) maximizes sensitivity, while a stricter consensus-of-five (logical set intersection) maximizes precision. We created five different annotation datasets by applying the consensus-of-*n* strategy to the experts’ annotations, from *n*=1 to *n*=5. Each consensus annotation dataset combined information from all five experts’ annotations using this thresholding scheme, and consisted of labels for the same set of independently annotated 20,099 images. Images for each Aβ class and each consensus strategy are shown in Figure 1C.

Qualitatively, we saw more phenotypic uniformity as more agreement was reached from *n*=1 to *n*=5. This held especially for the diffuse class in which some *n*=1 images resembled the phenotype for cored plaques, but as we increased to *n*=5, the classical phenotype emerged of sparse and scattered Aβ protein. A complete consensus-of-five experts was reached for 65% of images labeled as a diffuse plaque from any expert (Figure 1D). For the CAA class, the classical ring structure around blood vessels emerged as *n* increased. Complete consensus-of-five was reached for 33% of images with any CAA annotation (Figure 1D). We did not specify CAA subtypes^28,29^. For the cored class however, there was no smooth qualitative progression, visually indicating more differences and idiosyncrasies in identification. Complete agreement on a positive label only occured for 11% of cored-plaque labeled images. There were many cases in which only one annotator identified a particular cored plaque, with this consensus-of-one scenario making up the majority (44%) of cored-plaque images (Figure 1D).

To assess patterns of inter-rater agreement, we calculated the Cohen’s kappa coefficient^30^ between every pair of experts (Figure 2). There was low average agreement for the diffuse class (kappa = 0.46±0.11) despite the fact that a complete consensus (*n*=5) for it was the most common case out of any consensus. Hence, for diffuse cases without complete agreement, experts varied greatly. For CAA cases, average inter-rater agreement was much higher (kappa = 0.76±0.10). Agreement for the cored cases was low (kappa = 0.50±0.08). No single expert was a clear outlier in annotation across all Aβ classes. NP2 differed most with the other annotators for the diffuse and CAA classes, and NP4 differed most for the cored class.

**Figure 2.**
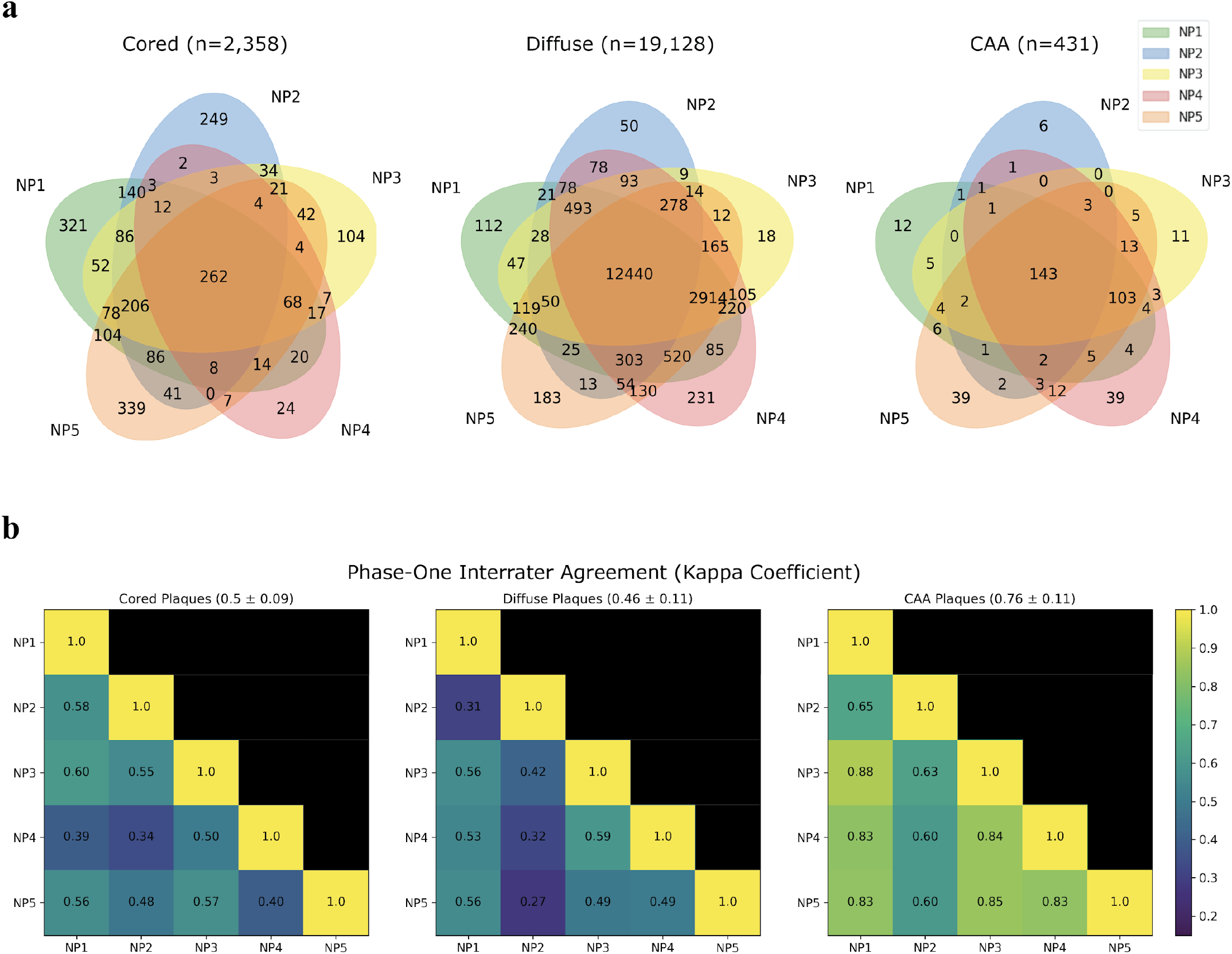
Inter-rater agreement varies by class and annotator. (a) Venn diagrams by class, with overlaps of each permutation of NP1 through NP5. Each overlap shows the count of how many images are all positively annotated by the experts included in that overlap. Areas are not to scale. (b) Kappa coefficients^30^ indicating agreement between each pair of experts. A high kappa coefficient indicates high inter-rater agreement between two annotators, with kappa = 1.0 indicating perfect agreement, and kappa = 0.0 indicating no agreement other than what would be expected by random chance.

### CNNs mimic human annotators, and consensus CNNs mimic a consensus of experts

CNNs are a powerful class of DL networks that are particularly useful for analyzing image data^31^. We found CNNs accurately learned the specific annotation behavior of both humans and consensus strategies. Using the five expert annotation datasets, we trained CNNs to reproduce each human’s annotations. We also trained independent CNNs to reproduce each consensus-of-*n* strategy.

We randomly assigned each WSI to either the training set or test set. Of the 20,099 center-cropped images, we held out 33% as a test set, and the remaining training images were split into a fourfold cross-validation (Methods). Both the models trained on individual-expert annotations and the models trained on the consensus annotations generalized and performed well on the held-out test set (Figure 3); they had high area under the receiver operating characteristic (AUROC) and area under the precision recall curve (AUPRC). We evaluated each model according to the labels of the annotation dataset on which it was trained. The consensus models mimicked consensus strategies, and the individual-expert models mimicked their corresponding expert annotators. Interestingly, the individual-expert models captured their human inter-rater agreement patterns, further suggesting that they were mimicking their human counterparts (Supplementary Figure 3). For every scoring metric and every Aβ class, consensus models performed slightly better than individual models (Figure 3). On average, they were able to more accurately reproduce the consensus strategies than the individual-expert models were able to reproduce their specific human annotators. There were substantial performance differences among the Aβ classes. All models were able to accurately reproduce the annotations of the diffuse class, which was far more ubiquitous than the cored and CAA classes. For these crucial but minority classes, AUROC and AUPRC performance was lower likely due to less available training examples (e.g., 1-2% for CAA class prevalence, depending on label strategy). Performance by stain was largely consistent, except for the CAA class with 6E10 staining, where performance decreased and was more variable (Supplementary Figure 4). Color-normalization had little effect on performance, except for the CAA class (Supplementary Figure 5).

**Figure 3.**
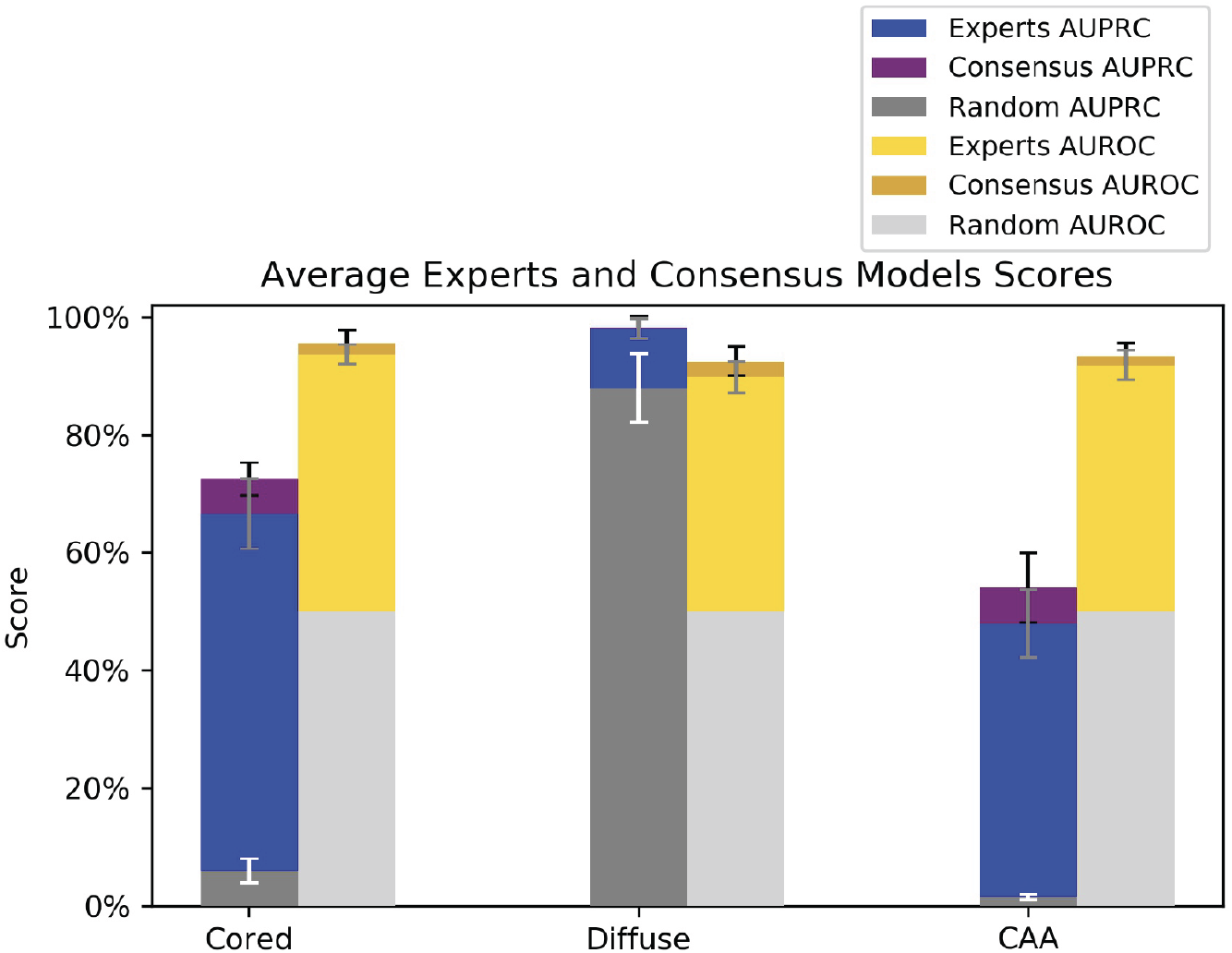
We trained models to learn human annotation behavior and consensus strategies. Consensus models matched or outperformed individual-expert models in average AUROC and AUPRC, per stacked bar graphs. Error bars show one standard deviation in each direction. The y-axis indicates the score on the held-out test set for each Aβ class (x-axis). No novice models were included in this evaluation. For the AUPRC metric, the consensus model achieved 0.73±0.03 for cored, 0.98±0.02 for diffuse, and 0.54±0.06 for CAA. The individual-expert models achieved 0.67±0.06 for cored, 0.98±0.02 for diffuse, and 0.48±0.06 for CAA. Random baseline performance for AUPRC is the average prevalence of positive examples. Average random baselines for individuals-experts were equivalent to those of consensus strategies (variance of individual-experts shown): 0.06±0.02 for cored, 0.88±0.06, and 0.02±0.004 for CAA. For the AUROC metric, the consensus models achieved 0.96±0.02 for cored, 0.92±0.02 for diffuse, and 0.93±0.02 for CAA. The individual-expert models achieved 0.94±0.02 for cored, 0.90±0.03 for diffuse, and 0.92±0.03 for CAA.

### Consensus provided superior models and annotation benchmarks

We found that learning from a consensus strategy resulted in consistently higher model performance than learning from individual-expert annotations. Since there was no clear best or established evaluation benchmark strategy across the test-set images (with each image having five independent expert annotations), we compared the consensus models with the individual-expert models using four different benchmark schemes (Figure 4A).

**Figure 4.**
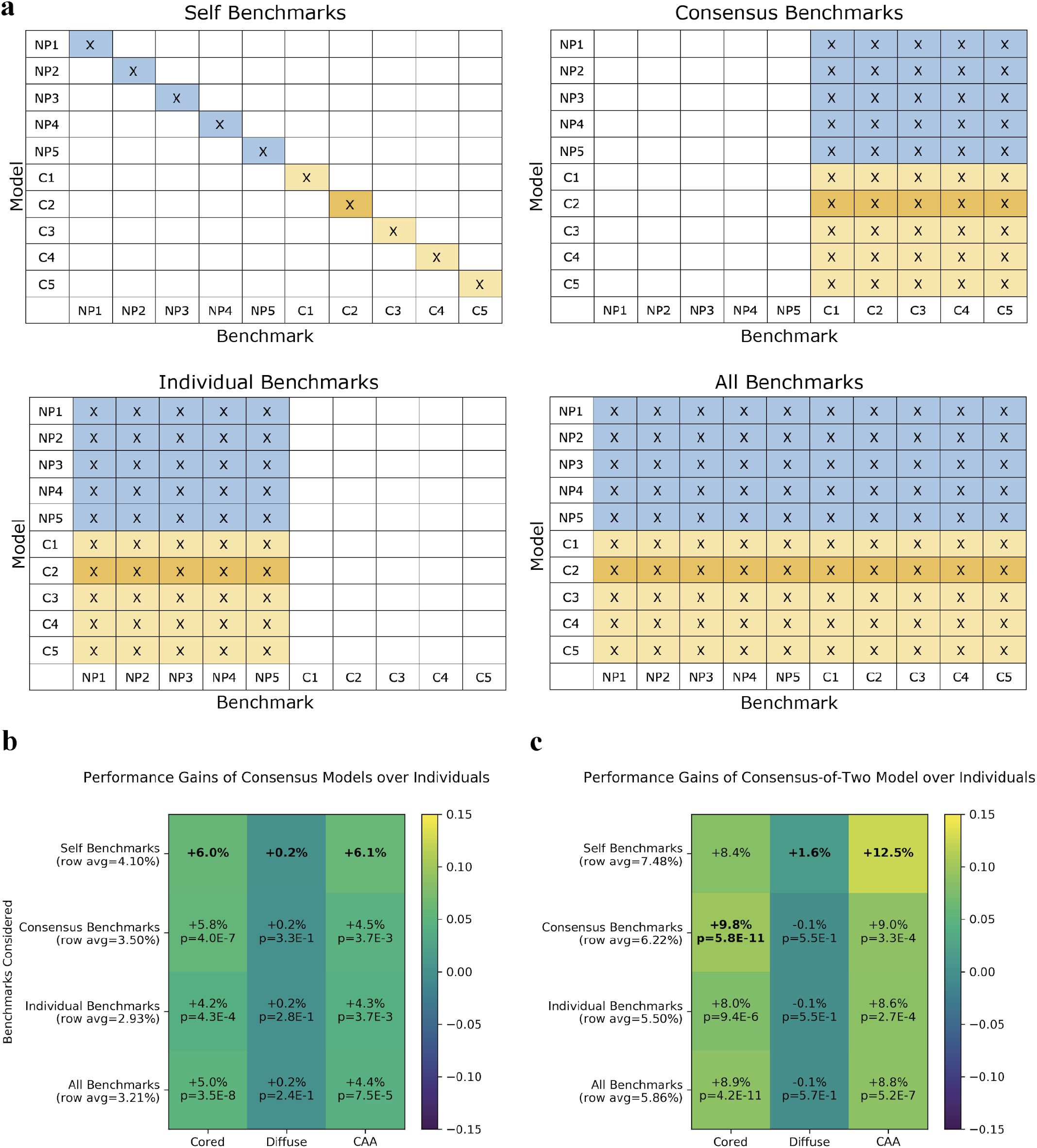
Consensus models performed better than individual-expert models across all benchmarks. (a) Four evaluation benchmark schemes to compare consensus models with individual-expert models. The row indicates the model and the column indicates the benchmark. For each evaluation scheme, the average AUPRC of the blue region (individual-expert models) is compared with the average AUPRC of the gold region (consensus models) over the held-out test set. The consensus-of-two is dark-gold for emphasis. The “self benchmarks” scheme was the most internally-consistent scheme that evaluated each individual-expert model according to the labels of its annotator (i.e. its own benchmark). For consensus models, the self benchmark corresponded to labels derived from the matching consensus-of-*n* strategy. The “consensus benchmarks” scheme independently evaluated each model on every consensus-of-*n* annotation set from *n*=1 to *n*=5. The “individual benchmarks” scheme independently evaluated each model on each of the five individual-expert benchmarks. The “all benchmarks” scheme evaluated each model on its average performance across all benchmarks. (b) Performance gains of consensus models over individual-expert models. Values are reported as the absolute AUPRC difference. We calculated p-values of the comparisons using a two-sample Z-test (Methods). P-values for the self-benchmark are not included because the sample size (*n*=20 comparisons) is not large enough to assign significance. The row indicates the type of benchmark considered when evaluating the model performance differentials, while the column shows the Aβ class being evaluated. Highest performance differential for each Aβ class in bold. (c) Heatmap as in (b), for only the consensus-of-two model versus the individual-expert models. For this consensus-of-two model evaluation, only dark-gold regions in (a) corresponding to the consensus-of-two model are compared to the blue region.

For all four benchmark schemes, the consensus models had superior average AUPRC performance versus individual-expert models (Figure 4B). We found this surprising, especially for the “individual benchmarks” case in which all models were evaluated with the different experts’ annotation benchmarks. We had expected individual-expert models to perform the best on this benchmark, because they were trained specifically to mimic these same annotators. Instead, we observed the consensus models—which in this scenario were trained under a different annotation dataset than the one that they were being evaluated with—consistently performed better on average than the individual-expert models across all Aβ classes (Figure 4B). Of the five consensus schemes, the consensus-of-two model achieved the highest average AUPRC (0.70±0.24) and AUROC (0.88±0.13) when evaluated against all benchmarks and all Aβ classes. On average, this consensus-of-two model substantially outperformed individual-expert models for all benchmark schemes and for all Aβ classes, except for the diffuse class in which performance was roughly equivalent (Figure 4C).

When we did the converse and compared annotation *benchmarks* as opposed to comparing *models*, we observed annotation datasets from consensus-of-*n* strategies provided a more robust benchmark yielding greater average model performance, regardless of which models we evaluated (Supplementary Figure 6A-B). Similarly, regardless of training strategy, models performed better on the consensus-of-two benchmark than on individual-expert benchmarks, for all Aβ classes (Supplementary Figure 6C).

### Model interpretation reflected differences in human expertise

Models learned human interpretable, pathologically relevant, and granular visual features for each Aβ class. By incorporating saliency mapping methods^25,32^ to interpret the model’s rationale, we found models focused on pixels corresponding to boundaries of Aβ pathologies and excluding boundaries of extraneous deposits in a way that was task (pathology) specific—even though this granularity of information was not provided during training (Figure 5A).

**Figure 5.**
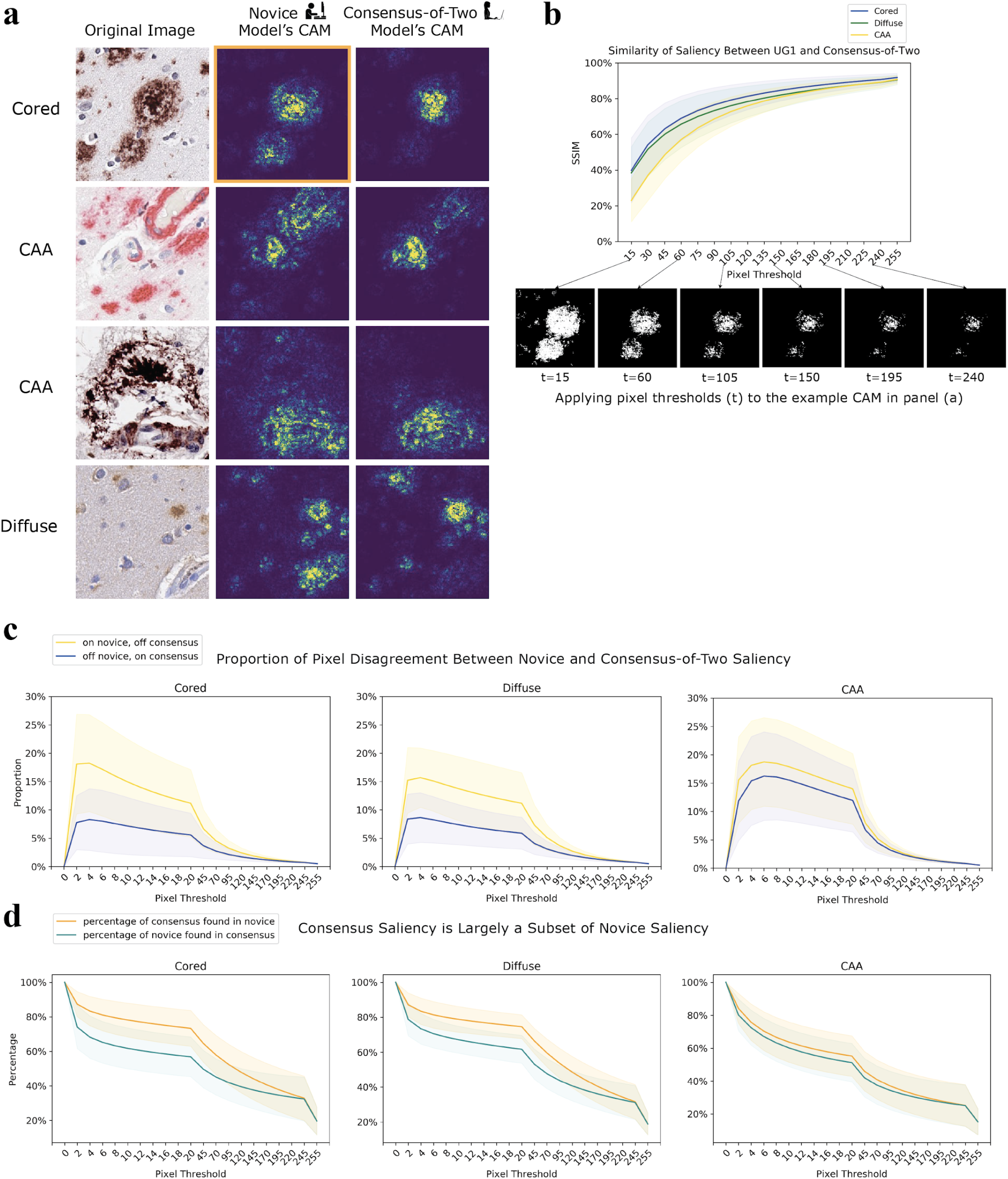
Class activation maps (CAM) of DL models indicate progression of human expertise. (a) Novice CAMs are more diffuse than expert CAMs. The original image (leftmost column), the CAM of the novice model trained on UG1’s annotations (middle column), and the CAM of the consensus-of-two model (rightmost column). CAMs are plotted with a false-color map such that bright regions correspond to high intensity regions with high salience. (b) Although expert and novice CAMs differ, they converge on the same pixels. We progressively assess the structural similarity index (SSIM)^41^ between novice CAMs and consensus-of-two CAMs across the entire test set of images. The CAMs show the most similar salience by SSIM (y-axis) at the highest pixel thresholds as we increment the threshold (x-axis) used to binarize the images before comparison. Binarized examples are shown of one CAM from (a) (boxed in orange). (c) Comparing the novice CAMs and the consensus-of-two CAMs, we classify each pixel location into two categories: ON in the novice CAM and OFF in the corresponding consensus CAM (yellow), or OFF in the novice CAM and ON in the consensus CAM (blue). ON and OFF are determined by binarizing the images at pixel threshold *t* (x-axis). Y-axis shows the proportions at which these two cases occur. Zoomed inset highlights disagreement between CAMs. (d) Consensus CAM pixels are mostly contained within the novice CAM. The x-axis plots the varying pixel thresholds, while the y-axis plots the percent overlap of either how much of the consensus CAM pixels are a subset of the novice CAM (orange) or how much those of the novice CAM are a subset of the consensus (cyan).

To investigate the impact of field expertise, we also trained models from the annotations of two undergraduate novices. We observed that a model trained on a consensus of two experts had more focused and specific feature saliency than a model trained on an individual undergraduate novice. To visualize what parts of an image were important to a model’s decision making, we used guided gradient-weighted class activation mapping, or “guided Grad-CAM”^25^ (Methods). When we compared class activation maps (CAMs) of the two models—consensus-of-two versus an undergraduate novice—we observed the CAMs of the novice’s model were more diffuse than the CAMs of the consensus-of-two model (Figure 5A). Qualitatively, the individual novice models attributed greater salience to pixel features that were not important to the classification task, while the consensus-of-two model focused on relevant morphologies. Therefore, the model trained on consensus across greater expertise appeared to be a more specific feature detector. These results held for both novice annotators (Supplementary Figure 7). However, subsequent studies with additional novices are needed to investigate this trend.

Quantitatively, for each Aβ class and for each image in our test set, we created CAMs and compared its novice CAMs to its consensus-of-two CAMs. For each pair of CAMs derived from the same image and the same class, but different model, we binarized the CAMs across an incrementing range of thresholds (Figure 5B). The majority of activations matched between the novice and consensus-of-two model at the different thresholds. In regions where they did not match, the novice CAMs tended to have signal while the consensus-of-two CAM did not (Figure 5C). Figure 5B shows binarized image examples at different thresholds. At high pixel thresholds the images became more similar as the granular features of the image were lost. Furthermore, we computed the fraction of the consensus-of-two CAMs found within the novice CAMs, and also the fraction of the novice CAMs found within the consensus-of-two CAMs across different pixel thresholds. The consensus-of-two CAMs were a smaller and more focused subset of the novice CAMs (Figure 5D).

### Ensemble learners bolstered performance and were robust to intentionally noisy “annotators”

We further tested multi-rater strategies by creating an ensemble of individual-expert models (no novices) and taking a “wisdom of crowds” approach. One approach would be to simply take the predictions of the different expert models and use a majority-voting scheme. However, weighting each model equally would ignore the specific and differing expertise the models gleaned during training. Other approaches assign differently weighted votes to different models^16^. Accordingly, we combined the individual-expert models in a learnable way, such that the resulting ensemble learned how to properly weight each contributing network’s Aβ class predictions to maximize overall performance. We theorized that the ensemble would learn how to combine the strengths of individual-expert models. The ensemble training did not modify the individual CNNs, but rather learned how to weight each constituent network to maximize performance for a given annotation benchmark (Figure 6A). We created ensembles for each of the phase-one annotation sets: both individual-expert annotation sets, as well as consensus annotation sets (Methods).

**Figure 6.**
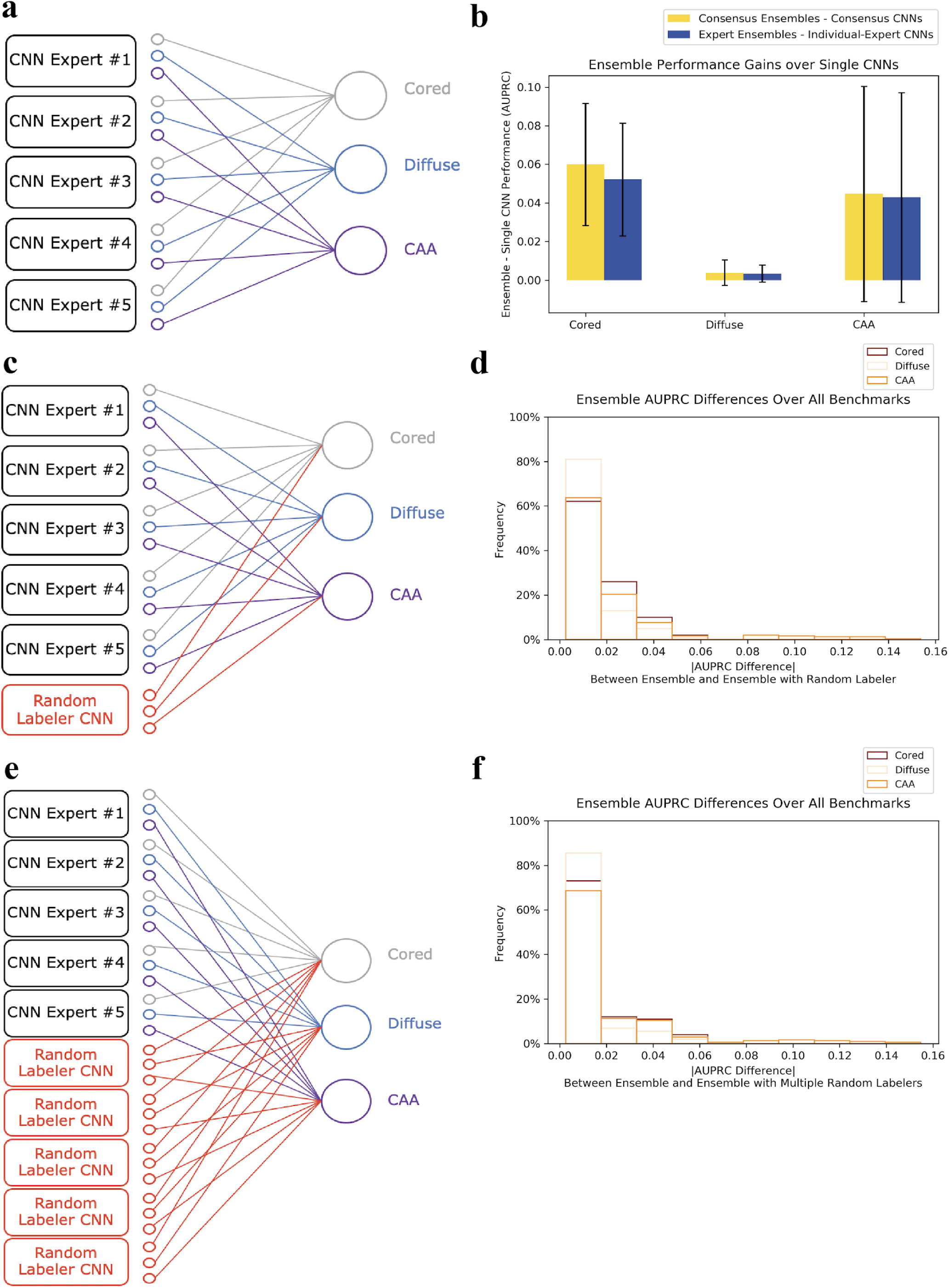
Ensembles improve performance and are robust to false information. (a) Five trained individual-expert CNNs, combined by a trainable sparse affine layer, make up an ensemble model. The training process simply determines how to best weigh and combine each CNN’s existing class predictions. (b) Ensembling on average increases performance for each Aβ class, and for both consensus and individual benchmarks. Performance gains are calculated by averaging each ensemble’s AUPRC on the held-out test set minus the correspondingindividual-expert CNN’s AUPRC on the same set, across all ten benchmarks (Methods). (c) We tested ensembling with a random labeler CNN, trained using a randomly shuffled permutation of labels with the same class distribution ratios as the five expert annotations. (d) Ensemble performance is largely unaffected by inclusion of a random labeler CNN. Density histogram of AUPRC performance differences for each Aβ class between the normal ensemble and the ensemble with a single random labeler CNN. Each ensemble is evaluated on all ten benchmarks (five individual-expert benchmarks, five consensus benchmarks), and the absolute value of the performance differential (x-axis) is calculated and binned for each class. (e) Ensemble architecture with multiple random labeler CNNs, each trained on a different permutation of randomly shuffled labels. (f) Ensemble performance is largely unaffected by inclusion of five random labeler CNNs. Same density histogram as in (d), but comparison is between normal ensemble and ensemble with five random labeler CNNs injected.

This ensemble approach bolstered performance, and improved over individual-expert CNNs in identifying CAAs. On average, ensembles outperformed single (non-ensembled) CNNs (Figure 6B). Performance differences for diffuse class detection were relatively unaffected, likely because these examples were already highly prevalent. Rather, we observed performance gains for the less prevalent but pathologically important minority classes (cored and CAA). For consensus ensemble cases, in which we trained different ensembles to reproduce consensus-of-*n* annotations from *n*=1 to *n*=5, the ensembles matched or slightly outperformed single CNNs trained to mirror the consensus.

Next, we investigated whether these performance gains were robust to having a poor annotator included in the ensembles (Figure 6C). We modeled this by creating a random labeler, who assigned randomly shuffled annotations that matched the average class distribution of all experts. When we included a random annotator CNN in the ensembles, the ensembles successfully learned to reject the noisy information from the random labeler. Performance was largely unaffected by the random annotator CNN, with differences in performance averaging to less than 0.01 in AUPRC across every permutation of ensemble model and evaluation benchmark (Figure 6D).

In a separate experiment, we also included five independent random labeler CNNs in the ensembles (Figure 6E), and we observed the same resilience and robustness. The ensembles successfully learned to ignore noisy contributors, and maintained performance even at a population that consisted of 50% poor labelers who inserted random information (Figure 6F). Hence, we found ensembles resulted in better performance, and were robust to multiple poor annotators at little perceived risk of random information. By contrast, including random annotations into the consensus-of-*n* schemes would proportionally dilute the training data signal.

### Annotators performed a second phase of annotation and were self-consistent

To assess use in a real-world clinical research setting, we designed a second, prospective, stage of the study (denoted as “phase-two”) to assess whether models could usefully serve as a prospective filter by identifying comparatively infrequent cored plaques and CAAs on new patients. Whereas cored plaques and CAAs have low prevalence (12% and 2% in phase-one, respectively), we posited that models from the phase-one could filter a large and previously unseen dataset of primarily diffuse plaques to enrich for these rare but important neuropathologies. We hypothesized these models could prospectively predict what their annotators were going to label. Moreover, given our initial findings that a consensus-of-two model was more effective and robust than individual-expert models, we aimed to compare their prospective capabilities.

We conducted phase-two annotations six months after the completion of phase-one. During phase-two, each annotator received a total of 10,511 images to label, using the same web platform as phase-one (Figure 7A). Annotators were told only that the study would proceed in two phases, but were given no information regarding phase-two image or data selection. Each blinded image came from one of four categories: “self-repeat,” “consensus-repeat,” “self-enrichment,” and “consensus-enrichment.” Self-repeats and consensus-repeats were collected to assess how consistently annotators were able to label images. These repeats were images the annotators had already seen in the first phase of annotation, and were triplicated, rotated randomly in 90-degree increments, and randomly spread throughout phase-two for intra-rater reliability testing (Methods). Self-repeats were random selections of images that the annotators labeled as positive during phase-one, balanced across all Aβ classes. Consensus-repeats were images with positive labels in the consensus-of-two annotation set from phase-one (Methods). Encouragingly, each annotator achieved high intra-rater agreement for all of the repeat images, and was able to consistently annotate and reproduce the same labels across the six-month gap (Figure 7B). Each expert achieved greater than 90% intra-rater accuracy among all three Aβ classes, while novices achieved greater than 88% accuracy. There was no significant difference in intra-rater agreement between self-repeats and consensus-repeats (Supplementary Figure 8).

**Figure 7.**
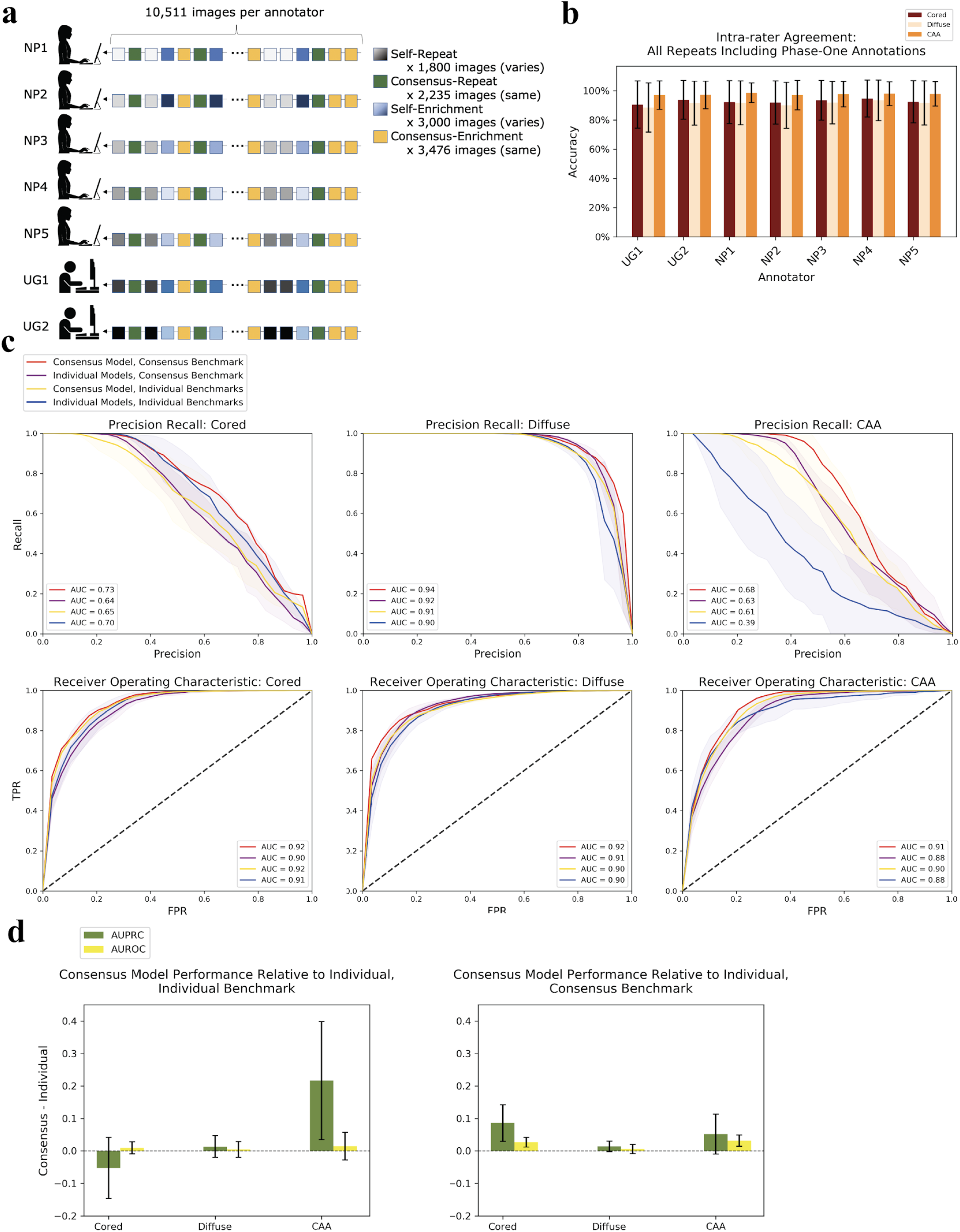
Models prospectively predict human annotation, with consensus models performing the most consistently. (a) Schematic of the phase-two annotation protocol. These images fall under one of four categories: self-repeat, consensus-repeat, self-enrichment, and consensus-enrichment. See Methods for a detailed description of these categories. Each annotator is given the same order of image categories. Gradients of different colors indicate images from the same category. These gradients are depicted to reinforce the fact that each annotator received a different set of images for the self-repeat and self-enrichment categories. (b) Intra-rater agreement is measured as the accuracy at which each rater consistently annotates repeats of the same image (both self-repeat and consensus-repeat). We include image labels from phase-one in this intra-rater calculation. The x-axis indicates the annotator, and the y-axis indicates intra-rater accuracy. Accuracies are averaged over each set of repeated images. Novices achieved an average intra-rater agreement accuracy of 0.92 for cored, 0.90 for diffuse, and 0.97 for CAA. Experts achieved an average intra-rater agreement accuracy of 0.93 for cored, 0.92 for diffuse, and 0.98 for CAA. (c) Precision recall plots and receiver operating characteristic (ROC) plots for the consensus model versus the individual-expert models. Two different benchmarks are used—truth according to the individual annotators, and truth according to a consensus-of-two scheme. The shaded regions indicate one standard deviation in each direction centered at the mean. The consensus model evaluated under a consensus benchmark (red line) has no variation by definition. (d) Summarizes panel (c). Bar graphs depict the average performance of the consensus model minus the average performance of the individual-expert models (y-axis). Individual benchmark for figure left, consensus benchmark for figure right. Error bars show one standard deviation centered at the mean.

Both self-enrichment and consensus-enrichment images were selected from unannotated and completely different WSIs than those used for phase-one, spanning 14 new patients across all three institutions. For the self-enrichment images assigned to annotator A, we used the individual-expert CNN trained on A’s phase-one annotations to enrich for images that the model predicted as having an important but minority plaque present (cored or CAA). For consensus-enrichment images, we used predictions from the consensus-of-two model instead to enrich for images. The same ordering of the four image categories was given to each annotator for consistency. Images differed between annotators in the self-repeat and self-enrichment categories only. See Methods for a detailed description of the different categories and selection process.

### Models prospectively enriched for minority Aβ classes and favored consensus training

Before completing this phase of the study, it was not clear whether a model trained on an individual annotator would prospectively perform the filtering and prediction task the best, as compared to a model trained from expert consensus. Consequently, from the 10,511 phase-two annotations across five expert annotators (i.e., total of 52,555 phase-two expert annotations), we created performance benchmarks to compare strategies. Models were assessed by either taking the individual expert’s labels as truth (called the “individual benchmarks,” five total benchmarks), or by taking a consensus-of-two experts as truth (called the “consensus benchmark”, one benchmark). We found that regardless of the selected benchmark, the consensus model performed better than individual-expert models in most cases, despite the fact that the individual-expert models were trained specifically under the same human annotators that provided the individual benchmarks (Figure 7C).

Both types of models were able to perform well prospectively for each Aβ class, with the consensus model outperforming individual-expert models on average in AUPRC and AUROC (by as much as 0.22±0.18 for CAA AUPRC, Figure 7C-D). For all classes and for all metrics, the consensus model evaluated on the consensus benchmark performed best. Furthermore, evaluating under a consensus benchmark as opposed to an individual benchmark resulted in either the same or better performance across models and across success metrics. Taken together the consensus model was robust to differing annotations of five different experts. On average, this model exceeded performance of individual-expert models even on their own benchmarks that they were trained to mimic, and also on the consensus benchmark (Figure 7D).

These results held for each class and each metric, with the sole exception being the cored class by AUPRC. In this case, individual-expert models slightly outperformed the consensus model under the individual benchmark. This was consistent with the phase-one observation that cored-plaque labeling had more idiosyncrasies (Figure 1C). Hence, we expected individual-expert models to be better performers on individual benchmarks for this class. Furthermore, evaluating individual-expert models under a consensus benchmark did not improve performance over the individual benchmark, likely for the same reason.

## Discussion

We developed a scalable, objective, and consistent way to automate annotations of neuropathology experts. Four points merit emphasis: 1) We automated individual and consensus annotations across datasets derived from five experts and three staining procedures from different institutions, advocating for generalizability; 2) When we prospectively validated these models in a research setting, a consensus model was robust under different benchmarks and provided a high-performance approach to automatic and standardized labeling; 3) The models were interpretable and showed increased pixel specificity with increased expertise; and 4) Ensemble models generally performed better and were robust to intentionally randomized information. We note however that the experimental setup and task differed from the daily practice of neuropathological annotation, where experts would commonly assess these objects using varying magnifications and localizations.

DL requires annotated datasets that define how to train a model and quantify its success. However, the classification of neuropathologies continues to evolve, with different experts having complementary focus areas. Consequently, two competing hypotheses regarding the best annotation strategy might be reasonable. In the first, a DL model trained on an individual might best leverage that single expert’s unique training, intuition, and decision-making procedures for edge cases. Conversely, in a second strategy, a consensus approach to annotations might instead leverage a wisdom-of-the-crowd logic to remove individual variance in a consolidated consensus model, as long as there is common underlying signal across the cohort. Depending on the consensus strategy, such a consolidated model could balance between strict-but-conservative expert consensus (the logical intersection) versus an inclusive but potentially overly-permissive consensus (the logical union).

In this study, we found that a balanced consensus-of-two strategy slightly favoring permissiveness performed better in nearly every model-training and model-benchmarking scenario (Figure 4C). This was striking, as the consensus-trained model outperformed the individual-expert models even when unfairly assessed against those self-same individual-expert benchmarks (Figure 4). One might reasonably have otherwise expected individual-expert models to better encode their particular expert’s pathological training and intuition, but this was generally not the case. We conclude from this observation that neuropathological labeling across thousands of independent annotations was sufficiently consistent across the cohort that a permissive consensus model computationally codified shared and commonly-held expertise in neuropathological identification—despite noticeable differences in individual annotations (Figure 2). We are unaware of previous work that operated on a cohort of independent expert annotations at this scale. Whereas other DL pathology applications rely on the independent assessments of individual pathologists^16,33,14,21,34^, we believe this is the first study to demonstrate that learning expert consensus within pathology provides robust and superior performance over learning individual assessments.

Encapsulating a consensus strategy into a model may be useful when an expert cohort or adequate labeling time are unavailable. For practitioners who would prefer individualized models or have unique research use-cases, the expert models effectively encoded individual intuition; thus they are limited in their application only by image-data preparation and computational-resource availability. Indeed, the purpose of phase-two was to prospectively assess whether these models could enrich for rarer but important neuropathologies within a large and unexplored dataset of n=275,879 candidate neuropathologies from new patients (Figure 7A). This represented a clinical scenario in which the pathologist wishes to rapidly curate sparse pathology examples for analysis, or to have a DL model act as an individualized assistant enriching for important but infrequent morphologies, which may improve understanding of these phenotypes and provide opportunity for overcoming sparsity challenges.

In clinical and research settings, we cannot rely on DL models solely as “black boxes”^35^ if we wish to have a common understanding of ground truth, guide diagnosis and other high-stakes decisions, or improve on subtle oversights of the current models. Leveraging the field of model interpretability, we thus employed a saliency mapping technique called guided grad-CAM^25^ to determine which pixels in the images contributed most strongly to classification. The models learned directly from the image data and became more focused in their feature detection as experience of the labeler increased from novice to expert consensus (Figure 5A). These results suggest potential for application in the training of new pathologists, using models to identify critical image features and thereby visually illustrate for new trainees the phenotypes relevant for accurate labeling. A future direction is to explore this new opportunity of DL facilitating human learning, as opposed to the more common framework of humans facilitating DL, ultimately leading to a virtuous hybrid feedback loop through active learning^36^.

The ensemble models may provide an opportunity of weighting individualized model input to tailor a specific annotator’s expertise to specific tasks or cases. Subsequent to a prediction, we can inspect the interplay of contributor-specific weights (Supplementary Figure 9). In this way, individualized contributions encoding complementary and situation-specific expertise do not get overwhelmed as they might otherwise under a simple vote averaging. Although we do not expect labelers to intentionally introduce false information, the ensembles’ robustness to unreliable annotations encourages assembling numerous and diverse labelers to facilitate accurate and learnable labeling. Scaling this approach to an even broader expert community remains a future direction.

Several limitations to this study inform its practical adoption. Researchers may not have access to necessary computational resources or training data for DL. However, cloud computing resources are becoming more affordable and accessible, and open-data sharing is also possible through data-hosting services. We are openly releasing this study’s annotated datasets and trained models (https://github.com/keiserlab/consensus-learning-paper). Turning to limitations in interpretability methods, we note that guided grad-CAM devolves to being a visual edge-detector in some settings^37^. However, this was not the case in our study because the saliency maps calculated on the same input image but different Aβ class substantially differed (Supplementary Figure 10). Nonetheless, a future direction would be exploring a range of independent saliency mapping techniques^38–40^. Finally, prospective effectiveness outside this cohort remains open to exploration. Whereas this study was consistent with our earlier work leveraging one expert annotator^14^ and its independent application at another institution^15^, we expect larger cohorts and institutionally-diverse datasets will facilitate more comprehensive standards in neuropathology. For instance, performance for CAA with 6E10 was lower than for other stains (Supplementary Figure 4). This could be due to its smaller representation in the test set (Supplementary Figure 11). Although this study’s cohort of five experts was institutionally diverse, it could be improved by capturing greater variability from the broader community. Despite these limitations, the models accurately learned the annotation practices of five experts and of their shared expertise, indicating the method’s generalizability.

We imagine this neuropathology study might apply as well to anatomic pathology and other areas of medical research. Any medical discipline that leverages human expertise could benefit from taking advantage of expert diversity and consensus to make accurate diagnoses. Models and annotated datasets developed and shared across the community could be progressively refined as ever-improving metrics and deployable tools of shared ground truth. Dissenting and novel views could likewise be codified, shared, and deployed to the community as functioning models and benchmark datasets. Whereas we focused on automatic image annotation, the concept may generalize to other domains and data types where expert annotation is crucial. Although a consensus-of-two experts was best in this particular study, this may not hold for other studies or for a different cohort or pathology, and indeed improved ensemble modeling techniques may be the strongest approach (Figure 6). However, the resilience and robustness of the consensus model indicated that training from a consensus was better than relying on individual assessments, despite high expert intra-rater reliability even over a half-year gap (Figure 7B). We hope that developing and openly sharing consistent, accurate, and automated DL methods and their datasets can facilitate standardization and accelerate quantitative pathology as a freely available community resource.

## Methods

### Slide Preparation

43 WSIs of the temporal cortex were collected from 3 different sites: 17 from UC Davis, 16 from University of Pittsburgh, and 10 from UT Southwestern. See Supplementary Figure 1 for associated clinical data. Slides were derived from formalin fixed paraffin embedded sections and stained with an antibody directed against Aβ. UC Davis used the Aβ 4G8 antibody, University of Pittsburgh used an NAB228 antibody, and UT Southwestern used a 6E10 antibody. All WSIs were imaged on an Aperio AT2 at 20X magnification.

### Data Collection and Annotation (Phase-one)

We selected 29 of the 43 WSIs at random for phase-one, and used the remaining 14 for phase-two. We color-normalized the WSIs according to the method presented in Reinhard^27^. Each WSI was uniformly tiled to 1536 x 1536 pixel non-overlapping images. After tiling, we applied a hue saturation value (HSV) filter and smoothing technique to detect candidate plaques, using the python library openCV. We used different HSV ranges for the different stain types as follows: 4G8 HSV = (0, 40), (10, 255), (0, 220); 6E10 HSV = (0, 40), (10, 255), (0, 220); cw HSV (0, 100), (1, 255), (0, 250).

Each candidate was center cropped to provide a 256 x 256 pixel image. This process yielded 526,531 images for phase-one. We randomly selected 20,099 images from this set to be annotated by our seven annotators. For the phase-two images, the center cropping yielded 275,879 images.

For the first phase of annotation, these 20,099 images were shuffled and placed into a fixed order, then uploaded to an Amazon instance web portal for independent annotation by seven different people. Five of them were professionally trained experts (E.B., B.N.D., M.E.F., J.K.K., K.E.M.), while the remaining two were undergraduate novices (J.A. and S.D.) with no formal training in neuropathology. They annotated the same 20,099 images in the same exact order in a multi-label classification task using a rapid-keystroke based custom annotation tool (first described in Tang et al.^14^, with current code released in https://github.com/keiserlab/consensus-learning-paper). Each 20X image had a bounding box, and the annotators were instructed to label any and all of Aβ pathologies found within the box. They had the option to label the pathology as any combination of three classes: cored, diffuse, or CAA. They also had the option to mark an image as negative, flag, or not sure (see Supplementary Figure 2). We did not use these alternative markings for constructing our final image labels. Hence, any Aβ class that was marked as positive was recorded as a positive annotation for that class. Any classes left unmarked were recorded as negative. There were instances in which an annotator marked negative and also marked a positive annotation for any of the three Aβ classes. Whenever this occurred (which was rare, Supplementary Figure 2B), we labeled the image as positive for the amyloid pathology indicated.

This process yielded seven different annotation sets (one for each person) for the 20,099 images that were annotated. From the five expert annotated sets, we constructed consensus-of-*n* annotation sets (from *n*=1 to *n*=5) for the same 20,099 images. For a given image *i* and Aβ class *c*, the new consensus-of-*n* annotation was recorded as positive if any *n* experts marked image *i* as positive for class *c*, else the image was recorded as negative.

### Training and Evaluation of DL Models

Of the 29 WSIs for phase-one, we used 20 WSIs for training and 9 separate WSIs for the held-out test set. The 20,099 phase-one images were divided into a 67% train and 33% held-out test split. Of the 67% training data, we performed fourfold cross-validation, keeping each fold’s image set consistent across all training and evaluation protocols such that each fold always had a distinct set of images that were not present in any other fold. We trained one model for each fold of the cross-validation, resulting in four models for each annotation set. Due to the large imbalance between the different Aβ classes, we performed class balancing during training. We calculated the ratios of diffuse to cored (*r1*), and diffuse to CAA (*r2*), and replicated any image with a cored plaque present a total of *r1* times, and any image with a CAA plaque present a total of *r2* times. Models were trained to perform multi-class classification of the three Aβ classes, taking a 256 x 256 pixel image as input and returning three floating point predictions (one for each class) as output.

For each of the twelve annotation sets (seven from our individual annotators, and five from our constructed consensus annotation sets), we trained four CNNs (one CNN for each fold). We trained models for 60 epochs, and saved the best model state that resulted in the highest cored AUPRC performance over the validation set. We used an Adam optimizer with a learning rate of 0.001, and a weight decay of 0.03. We used a multi-label soft margin loss as our loss function. During training, the input images were transformed with a random horizontal flip, a random vertical flip, a random 180 degree rotation, a color jitter, and a random affine transform so that the model would learn to generalize across different inputs. Images were normalized to have zero-mean and unit-variance prior to entering the model.

We converted each image annotation into a floating point representation, such that the binary labels given by the annotators were converted to a more specific continuous value, using information from other annotated bounding boxes that may be present in the image. During the annotation process, each 20X image possibly contained regions that were previously annotated in a different query with a different bounding box. Hence we had instances in which parts of bounding boxes or multiple bounding boxes from different annotations (from the same annotator) were present in a single 20X image. For each image *i*, we used the labeled bounding box information that the annotator provided in order to create a floating point representation for each class *c*. The floating point representation of *c* was the total sum of fractions of positively labeled bounding boxes for class *c* that exists within *i.* An image potentially had more than one positively labeled bounding box within *i,* or many fractions of positively labeled bounding boxes within *i.* Therefore, the floating point representation was possibly greater than 1.0, but must have been greater than or equal to 0.0. Finally, image *i* was considered positive for class *c* if the floating point representation of class *c* was greater than 0.99. This final binary label was used for training and evaluation.

During evaluation of a model, no transforms were applied except for normalization. For each type of model, we evaluated our models by taking each model from each fold and evaluating it on the held-out test set. The reported metrics were the average over these four evaluations. For example, the reported phase-one results for the consensus-of-two model was the average performance of all four consensus-of-two models, one for each of the four cross-validation folds, each evaluated on the held-out test set. The full source code used for training and evaluation can be found at https://github.com/keiserlab/consensus-learning-paper.

### Assessing Consensus Model and Benchmark Superiority

When assessing the consensus models versus the individual-expert models, we applied four different benchmark schemes using only the held-out test set annotations, and calculated the average performance of the consensus models minus the average performance of the five individual-expert models (Figure 4). “Self” means that each model was evaluated by its own benchmark (i.e. a model trained under A’s annotation set is evaluated according to the annotations in A’s held-out test set). For instance, a consensus-of-two model with benchmark “Self” was evaluated according to how well the predictions matched with the held-out test labels provided by the consensus-of-two annotations. This was done for each of the four consensus-of-two models (one for each fold), and averaged (Figure 4B). “Consensus benchmarks” means that all models were evaluated on how well their predictions matched on average with each of the five consensus benchmarks (consensus-of-*n* from *n*=1 to *n*=5). We averaged performance across these five consensus benchmarks. “Individual benchmarks” means that all models were evaluated on how well their predictions matched on average with each expert benchmark. We averaged performance across these five expert benchmarks. “All benchmarks” is evaluating a model across the five consensus benchmarks and five expert benchmarks, and then averaging these results. For the consensus-of-two superiority results (Figure 4C), we performed the same procedure, except only the consensus-of-two model and consensus-of-two benchmark were evaluated to represent the consensus strategy.

We used a two-sample, one-sided Z-test to assess statistical significance. The null hypothesis was that the average of the consensus performance and the average of the individual experts’ performance were the same. The alternative hypothesis was that the average of the consensus performance was greater than the average of the individual experts’ performance. P-values were reported using a standard lookup table. The sample sizes for each of the two samples (consensus and experts) are as follows: 20 for “self”; 100 for “consensus benchmarks”; 100 for “individual benchmarks”; and 200 for “all benchmarks”. Each sample was a model’s performance on a specific benchmark.

### Model Interpretability

We used guided Grad-CAM to obtain the saliency maps from the different models. The library we used can be found here: https://github.com/utkuozbulak/pytorch-cnn-visualizations. For analyses requiring image binarization, we chose incremental pixel thresholds such that at threshold *t*, any signal less than *t* is assigned 0 value, while anything greater than or equal to *t* is assigned a maximal pixel value of 255. To calculate SSIM between the novice CAM and the consensus-of-two CAM, we used a standard SSIM function from the skimage library.

For analyzing the CAM patterns of the novice CAMs versus the consensus-of-two CAMs, we constructed a subtraction map of the binary consensus CAM minus the binary novice CAM for each 256 x 256 pixel image. Each pixel of this map is classified as one of three classes: signal (ON) in the novice CAM and no signal (OFF) in the consensus CAM, or no signal in the novice CAM and signal in the consensus CAM, or a match between the two. For the CAM subset analysis (Figure 5D), we calculated both the total fraction of the binary consensus CAM that was activated in the binary novice CAM at the same corresponding pixel locations, and also the total fraction of the novice CAM that was activated in the consensus CAM at the same corresponding pixel locations for each pixel threshold.

### Ensemble Training and Evaluation

For each of the four folds of the cross-validation, we linked each of the five individual-expert CNNs trained from that fold with a sparse affine layer. The individual-expert CNNs were frozen such that no backpropagation occured in the individual-expert CNNs. The only weights that were updated were the spare affine weights, which weight each individual-expert CNN’s final class output.

For each annotation set (five expert sets, and five consensus annotation sets), four ensembles (one for each cross-validation) were trained using the same training data that their constituent CNNs used for training. Holistically, having four cross-validation folds, five experts, and five consensus schemes, resulted in 40 ensemble models, each independently trained to reproduce a specific annotation set. Ensemble training occurred for 60 epochs, with the same hyperparameters that were used to train the single CNNs.

After training, each ensemble model was evaluated on the images from the held-out test set, and on every benchmark (five expert benchmarks, and five consensus benchmarks). We aggregated ensembles that were trained to mirror the same annotation set but belonged to different cross-validation folds. Hence, every permutation of 1) ensemble model (aggregated across cross-validation folds) and 2) benchmark was assigned an average AUPRC score. This resulted in 100 average AUPRC scores (10 aggregated ensembles, 10 held-out test benchmarks). To assess ensemble superiority over single CNNs for a given benchmark, these 100 average AUPRC scores were compared to the 100 average AUPRC scores of the single (non-ensemble) CNNs. This resulted in 100 comparisons of ensemble models versus single CNNs, with both model types evaluated on equivalent benchmarks and an equivalent image set (Figure 6B).

For ensembles that contained a random labeler CNN, we trained a separate CNN on a random annotation set that kept the same class ratios of cored, diffuse, and CAA as the ones present among the five expert annotations. This ratio was determined by averaging the class ratios of each of the five expert annotation sets. We then linked this CNN trained on random labels with the five professional CNNs using a learnable sparse affine layer. Likewise, for ensembles with five random labeler CNNs, we trained five independent CNNs on five different permutations of the randomly labeled annotation set. We then linked these five random labeler CNNs with the five professional CNNs using a sparse affine layer. For both the ensemble with a single random labeler, and the ensembles with multiple random labelers, we used the same training procedure as the normal ensembles (i.e. ensembles without any random labeler present).

In evaluating the performance of ensembles with any number of random labeler(s) present, we performed the same evaluation procedure as for the normal ensembles. For comparing performance between ensembles with any random labeler versus performance of normal ensembles (Figure 6D, 6F), we compared the final average AUPRC values for these two model types and for each of the ten benchmarks. This resulted in 100 comparisons between normal ensembles and ensembles with a random labeler, and 100 comparisons between normal ensembles and ensembles with five random labelers. In every comparison, everything was kept constant except for the choice of model architecture.

### Setup of Phase-Two Prospective Validation and Analysis

For phase-two, we used 14 WSIs that were completely separate from the 29 WSIs from phase-one. We performed a second round of human annotation (phase-two) to validate the models and assess intra-rater reliability. All annotations were performed on the same web interface and by the same expert and novice annotators from phase-one. Annotators saw four different categories of images: self-repeat, consensensus-repeat, self-enrichment, and consensus-enrichment. For the self-repeat images, which are simply repeats of a subset images the annotators already saw in phase-one, we selected a total of 600 images. We first selected all of the images that were marked as positive for CAA. CAA was rarely annotated during phase-one, and we wanted to include all of them. Once all of the CAAs were included, we selected a random subset of images that were marked as positive for cored until we had 400 images. If there weren’t enough cored positive images to make up 400, then we took as many as possible. For the remaining images we randomly selected diffuse positive images to make up a total of 600 images. We enforced having no duplicates. We then triplicated and randomly rotated these 600 images, and shuffled the resulting 1,800 images. This entire procedure of obtaining the self-enrichment image set was done independently for each expert and each undergraduate novice, resulting in different sets of self-repeat images given to different annotators.

Likewise for the consensus-repeat set, we took images exclusively from the set of phase-one images that the annotators already labeled. For each of the three Aβ classes, we randomly selected 250 images that were positive for this class according to a consensus-of-two strategy. From this list of images, we removed duplicates, which occurred because some images were positive for multiple Aβ classes according to a consensus-of-two. This resulted in a total of 745 images, which were then triplicated and randomly shuffled, resulting in our final consensus-repeat set of 2,235 images. This set was given identically to all of the annotators during phase-two.

For the self-enrichment set given to annotator A, we used the CNN trained on individual A’s annotation set to select images that the model predicted as having a minority Aβ plaque present. We chose to use the models from fold three of the cross-validation for enrichment. All images came from a held-out set of images that the annotators did not see during phase-one of annotation. This held-out set consisted of 275,880 images total. From this held-out set, we randomly selected 800 images with a model prediction threshold > 0.90 for the cored class. Next, the 275,880 images were sorted according to the model’s CAA prediction confidence, and the top 800 images with highest CAA confidence were included. After collecting this set of 1,600 images, we included all of the image neighbors that had at least a 20% bounding box overlap of a plaque with any of these 1,600 images. Afterwards, we randomly shuffled this resulting list, and took a random subset of 3,000 images to use for our final self-enrichment set. This procedure was repeated for each of our seven human annotators. We calculated the overall intra-rater agreement for each annotator (Figure 7B) by averaging the accuracy of how consistent each annotator was over each set of replicated images. Each replicated set had four total images (one annotation from phase-one, and three annotations from phase-two).

For the consensus-enrichment set, we randomly pulled 750 images that the consensus-of-two model predicted as positive for cored, and an additional 750 images that the same model predicted as positive for CAA. From this resulting set of 1,500 images, we found their image neighbors that had at least a 20% bounding box overlap of a plaque with any of the 1,500 images. We randomly selected from these neighbors until we had a total of 3,476 images for the final consensus-enrichment set. If an image achieved high rank by both self-enrichment and consensus-enrichment, it was assigned with equal probability to either self or consensus.

These four image sets were given to the annotators, such that the ordering of the image category was randomly shuffled, but fixed and identical among each annotator. Upon completion of annotation, we analyzed the model’s prospective performance to match annotations given during phase-two. We stipulated two different benchmarks that could be derived from these new annotations: the individual benchmark, which simply assigned what the individual annotator labeled as the truth labels, and the consensus benchmark, which used the consensus-of-two strategy to assign truth labels. We used the individual CNN models to make class predictions for the 10,511 images of phase-two, independent of the human annotators and their labels. We did the same for the consensus-of-two model, and made class predictions for each of the 10,511 images. Model predictions were completely hidden from the annotators, as well as any experimental details of how the images were selected. For analysis on phase-two data, we define an individual-expert model as a model trained on one of the expert’s annotation sets from phase-one, and a consensus model as the model trained on the consensus-of-two annotation set from phase-one. All label data used to assess model performance during phase-two came from this second phase of annotation, and we did not use any phase-one annotation labels.

To assess performance of individual-expert models under the individual benchmark (shown in blue in Figure 7C), we compared the predictions of each individual-expert model trained on annotator A with the labels that annotator A gave during phase-two (undergraduate novices were excluded from this analysis). This was repeated for each expert annotator model and the five results were averaged. To assess performance of individual-expert models under a consensus-of-two benchmark (shown in purple in Figure 7C), we compared the individual-expert model’s predictions with the labels provided by a consensus-of-two scheme. This was done for each professional annotator model and the five results were averaged. For the consensus model and individual benchmark case (shown in yellow in Figure 7C), we compared the consensus model’s predictions with the labels provided by annotator A. This was repeated for each of the five professional annotator labels, and these five results were averaged. For the consensus model and consensus benchmark case (shown in red in Figure 7C), we compared the consensus model’s predictions with the label set provided by a consensus-of-two strategy. There was only one consensus-of-two model and one consensus benchmark, resulting in no variability and no averaging in this case.

## Data Availability

The raw WSIs, the color-normalized 1536 x 1536 pixel tiles, and the final 256 x 256 pixel images are all freely available, and can be found at: https://osf.io/xh2jd/

## Code Availability

All code and models are freely available at: https://github.com/keiserlab/consensus-learning-paper

## Acknowledgements

The authors thank the families and participants of the University of California Davis, University of Texas Southwestern, and the University of Pittsburgh Alzheimer’s Disease Research Centers (ADRC) for their generous donations as well as the ADRC staff and faculty for their contributions. This work was supported by grants from the National Institutes of Health: AG062517 (B.N.D.), University of California Office of the President (MRI-19-599956, B.N.D.), grant number 2018-191905 from the Chan Zuckerberg Initiative DAF, an advised fund of the Silicon Valley Community Foundation (M.J.K.), the California Department of Public Health (19-10611 B.N.D.), AG010129 (UC-Davis Alzheimer’s Disease Research Center), P30 AG013854 (M.E.F.), K08AG065463 (M.E.F.), P30 AG066468 (ADRC, J.K.K.), P50 AG005133 (ADRC, J.K.K.), P30 AG012300, the McCune Foundation, and the Winspear Family Center for Research on the Neuropathology of Alzheimer Disease (C.L.W.). We thank the UCD Health Department of Pathology and Laboratory Medicine for the use of their digital slide scanner.

## Author Contributions

Conceptualization, D.R.W., Z.T., B.N.D., and M.J.K.; Methodology, D.R.W., Z.T., B.N.D., and M.J.K; Software, D.R.W., Z.T.; Validation D.R.W.; Formal Analysis, D.R.W; Investigation, D.R.W., Z.T., S.D., J.A., K.E.M., J.K.K., M.F., E.B., B.N.D.; Resources, K.E.M., J.K.K., M.F., E.B., C.L.W., A.B., B.N.D., and M.J.K.; Data Curation, D.R.W, Z.T., N.M.; Writing – Original Draft, D.R.W.; Writing – Review & Editing, D.R.W., B.N.D., and M.J.K; Visualization, D.R.W.; Supervision, B.N.D., and M.J.K.; Project Administration, D.R.W., B.N.D., and M.J.K.; Funding Acquisition, B.N.D., and M.J.K.

## Declaration of Interests

The authors declare no competing interests.

## Notes

https://github.com/keiserlab/consensus-learning-paper

https://osf.io/xh2jd/

### Competing Interest Statement

The authors have declared no competing interest.

