## Supplementary Information for "Deep learning from multiple experts improves identification of amyloid neuropathologies"

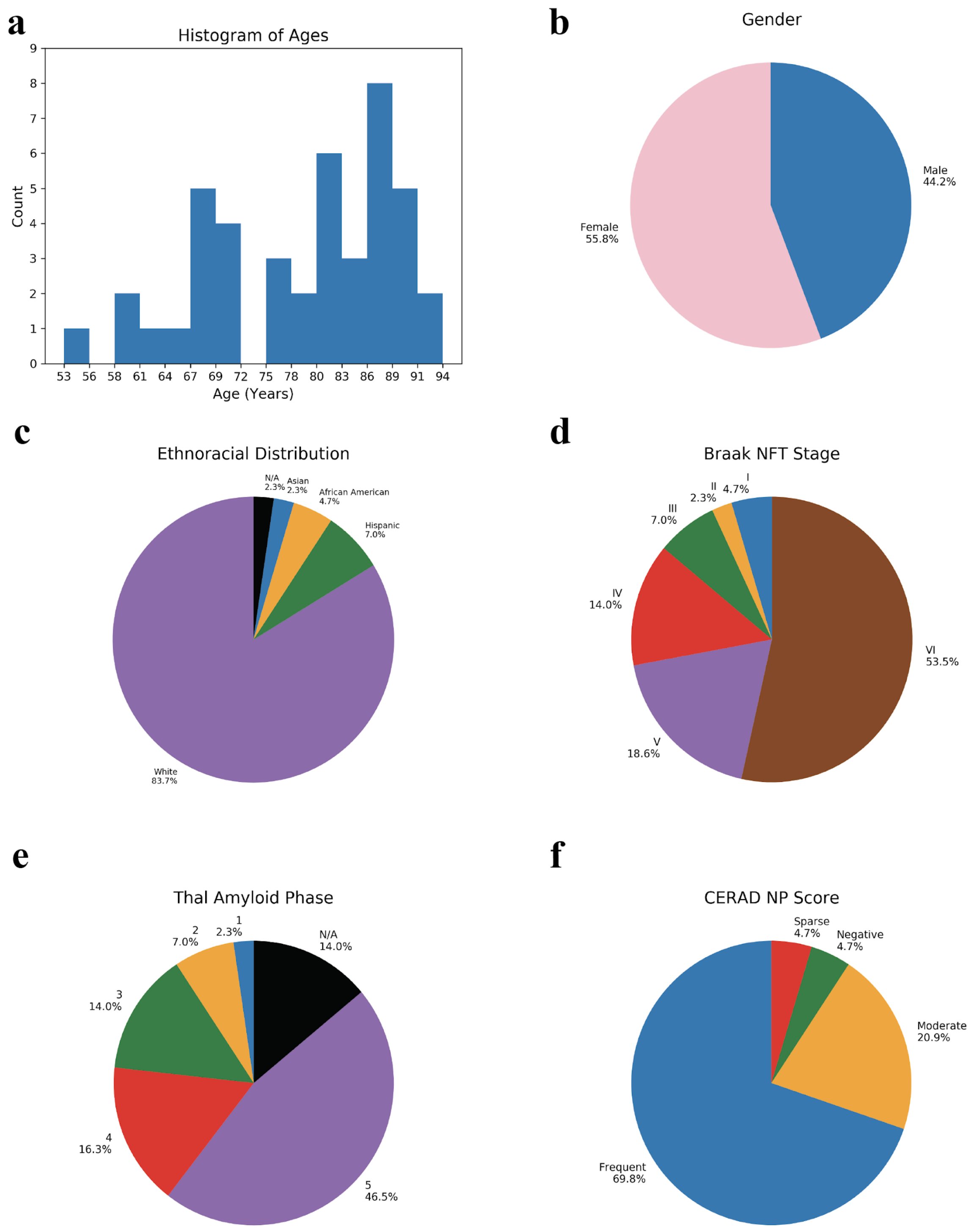


**Supplementary Figure 1. Demographics of the 43 patients from all institutions.** (a) Histogram of ages (b) Pie chart of gender distribution (c) Pie chart of racial distribution (d) Pie chart for scoring guidelines for Braak NFT Stage (e) Thal Amyloid Phase and (f) CERAD NP Score, set forth by Montine et al[1].

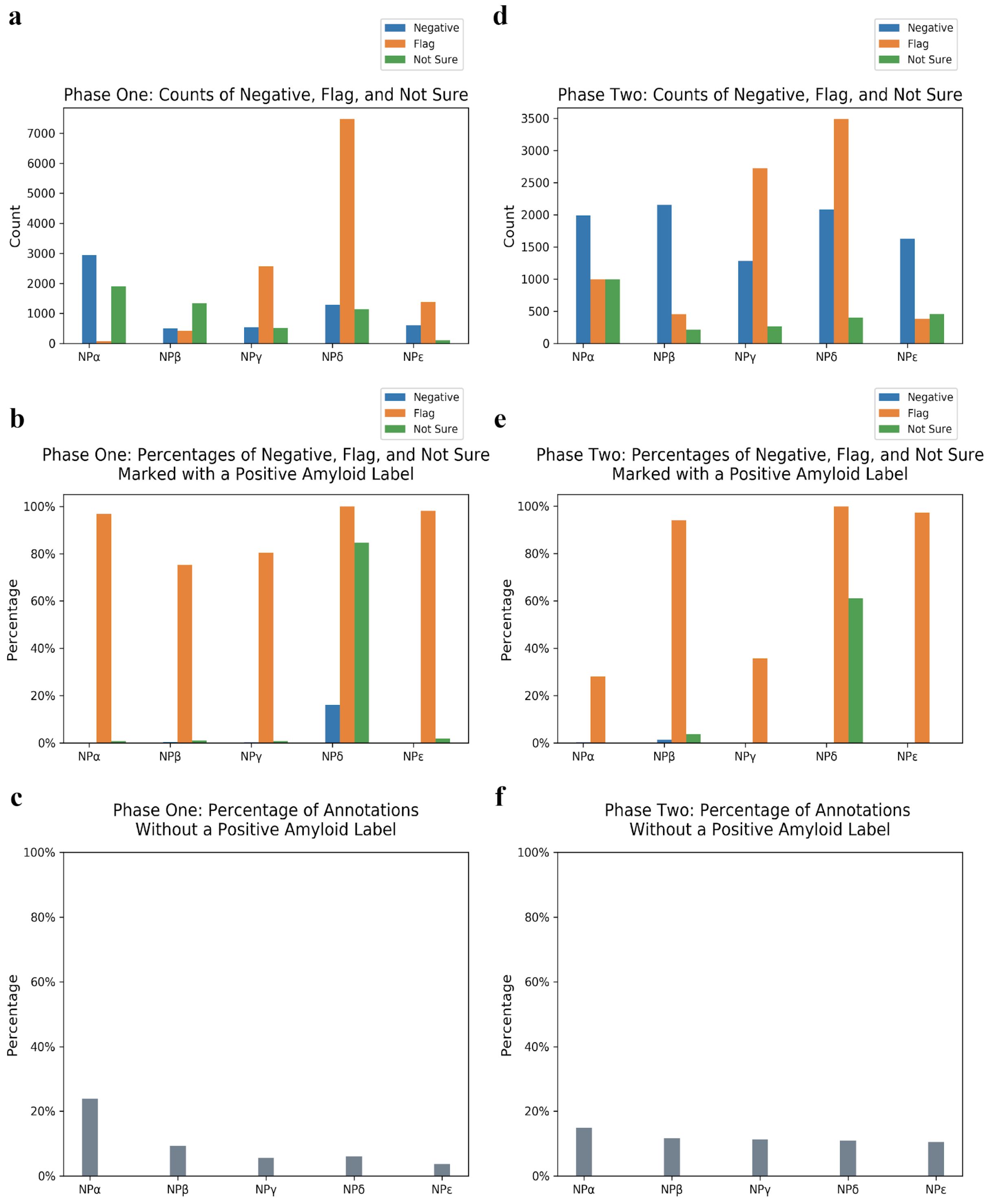


**Supplementary Figure 2: Negative, flag, and not sure annotations for both phase one and phase two.** For all panels, we further obfuscate the neuropathologist identities by assigning each with a randomly chosen but fixed Greek letter. (a) The number of times that an annotator checked one of the three alternatives: “negative”, “flag”, and “not sure” for phase one. There are a total of 20,099 annotations for each neuropathologist during phase one. (b) Of the three alternatives, the fraction of annotations in which the annotator also marked any of the amyloid plaques as being present during phase one. The cases in which annotators marked “negative” and also identified at least one amyloid plaque were sparse. In such erroneous cases, we chose to accept the positive amyloid marking(s) over the “negative” marking. For the majority of cases in which annotators marked “flag,” the annotators also identified at least one amyloid plaque as well. For these cases, we constructed the final image labels by using the positive amyloid marking(s), essentially disregarding the “flag” marking. For the cases in which an annotator marked “not sure” and also identified at least one amyloid plaque as well, we likewise used the positive amyloid marking for our final image labels. (c) Fraction of phase one annotations that did not receive any positive label for any of the three amyloid classes. (d) Counts of the three special cases for phase two. There were a total of 10,511 images. (e) Proportions of the three special cases that were also marked with a positive amyloid label during phase two. (f) Fraction of the phase two annotations without a positive amyloid label for any class.

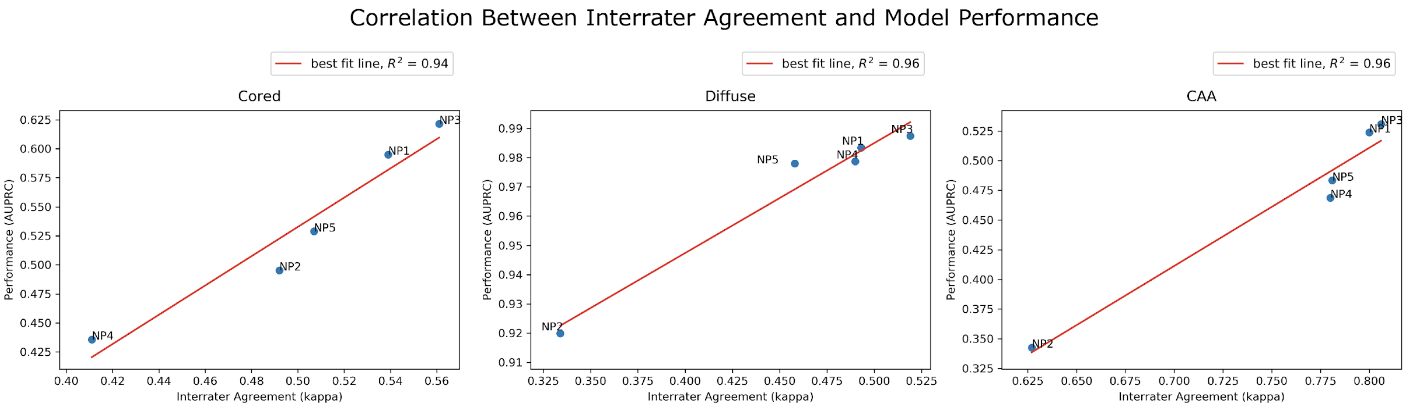


**Supplementary Figure 3. Models capture interrater agreement patterns among experts.** For each expert *E*, we calculated an average kappa coefficient between *E* and every other annotator. We also calculated the average AUPRC performance for expert *E*. We took every other expert model that was not trained with *E*'s annotation set, and evaluated those models using *E*'s annotation benchmark, and averaged the results. This yielded a single kappa score (x-axis) and a single AUPRC score (y-axis) for each expert *E*. Correlating these two scores, we saw human inter-rater agreement reflected in the ML models. We calculated $R^2$ correlation from the interpolated line of best fit. There were strong $R^2$ correlations (0.94 for cored, 0.96 for diffuse, and 0.96 for CAA) between an individual's kappa coefficient and how well other models acting as annotators agreed with this individual's benchmark.

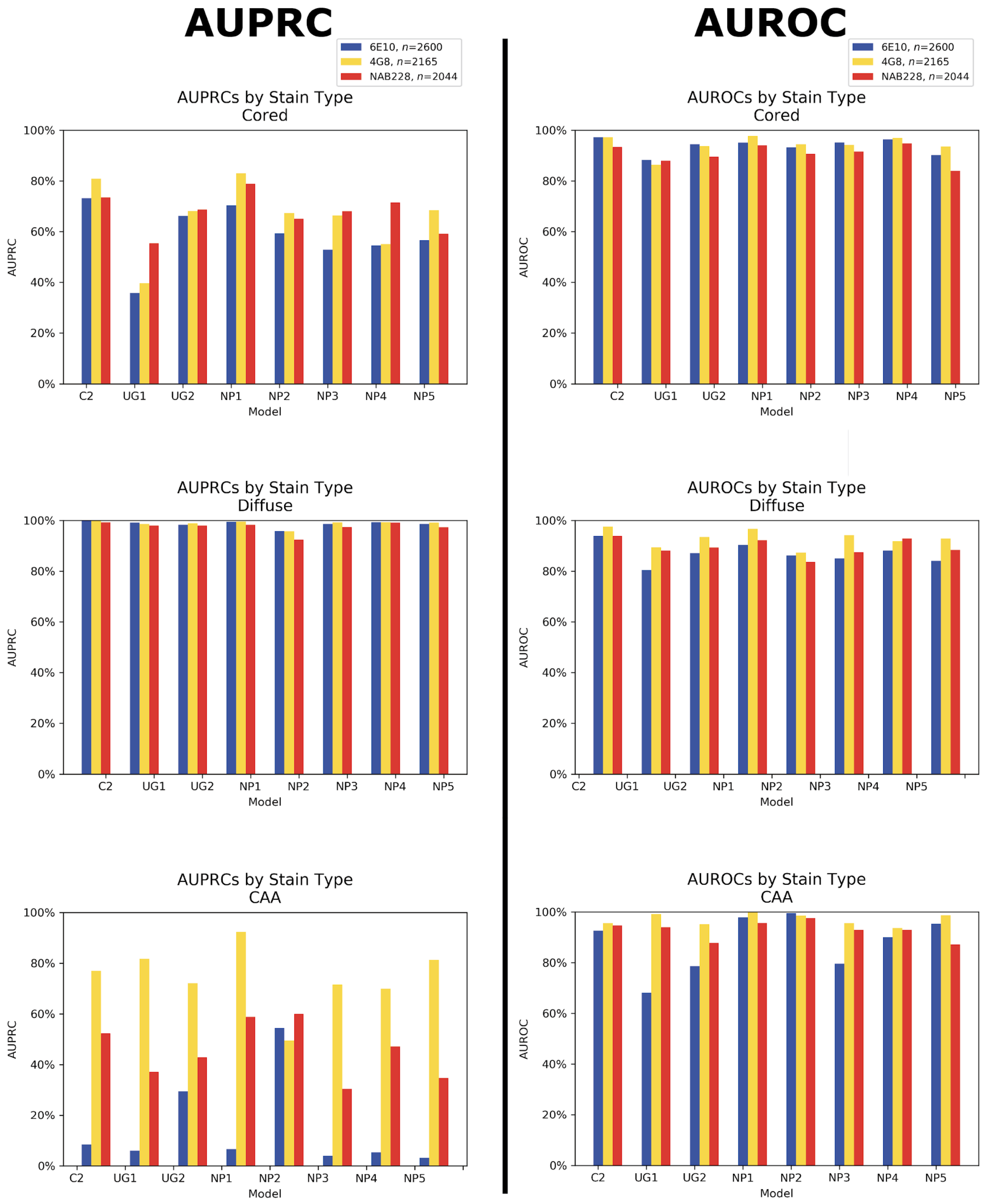


**Supplementary Figure 4: Model performance by stain type for the three Aβ classes.** The left half of the figure is AUPRC, while the right half is the corresponding AUROC. The x-axis designates the model. "C2" refers to the consensus-of-two model. The y-axis designates the performance metric over the held-out test set. There are three different stain types, each from three different institutions: 6E10, 4G8, and NAB228. The number of images (*n*) per stain type in the test set are displayed in the figure legend. Both AUPRC and AUROC performance are consistent across stain types. The exception is AUPRC for CAA with the 6E10 stain. In this case, representation in the test set is small (Supplementary Figure 11). Since each stain came from a different institution, effects from stain versus patient cohort cannot be disentangled.

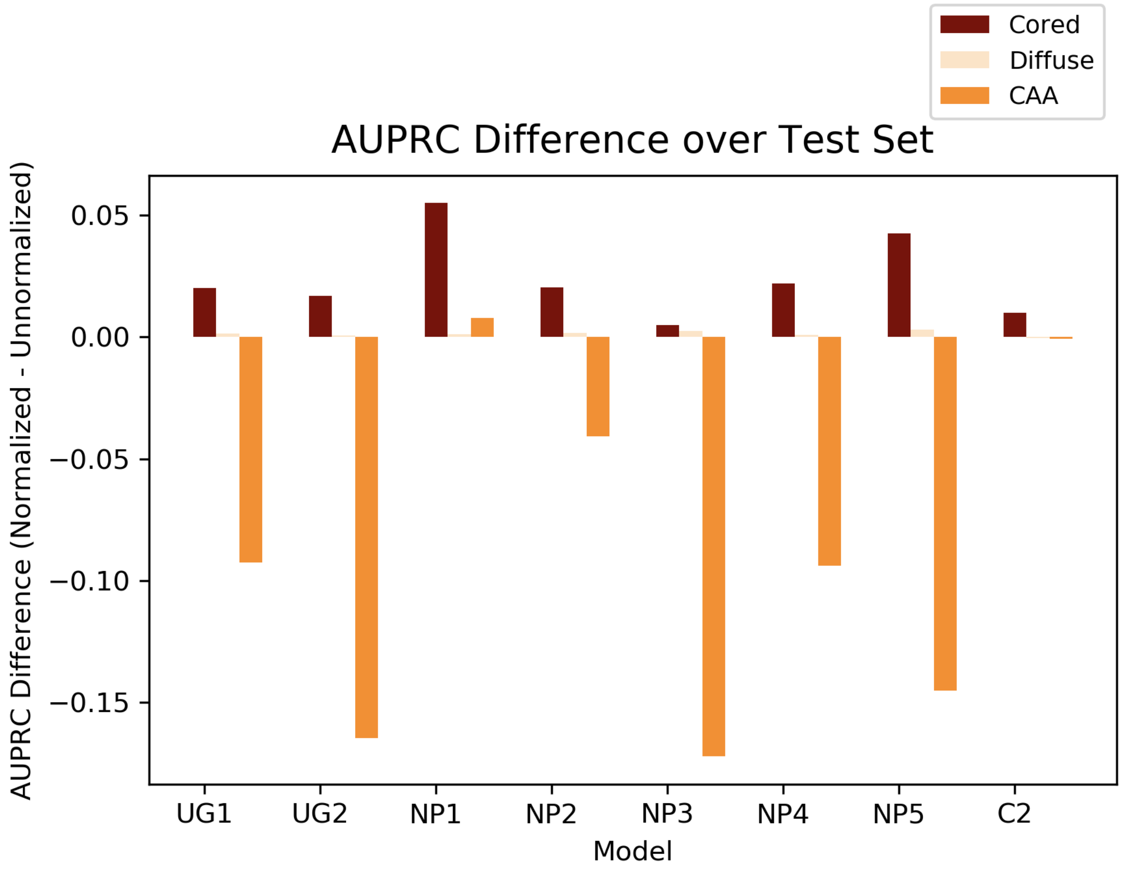


**Supplementary Figure 5: Model evaluation differences on color-normalized versus unnormalized test images.** Each model (x-axis) is evaluated on its own benchmark's test set. "C2" is the consensus-of-two model. The difference in AUPRC (y-axis) is calculated when evaluating the model on color normalized images versus unnormalized images. Results are stratified by amyloid class. Most models have higher performance for the CAA class when we do not color-normalize the images.

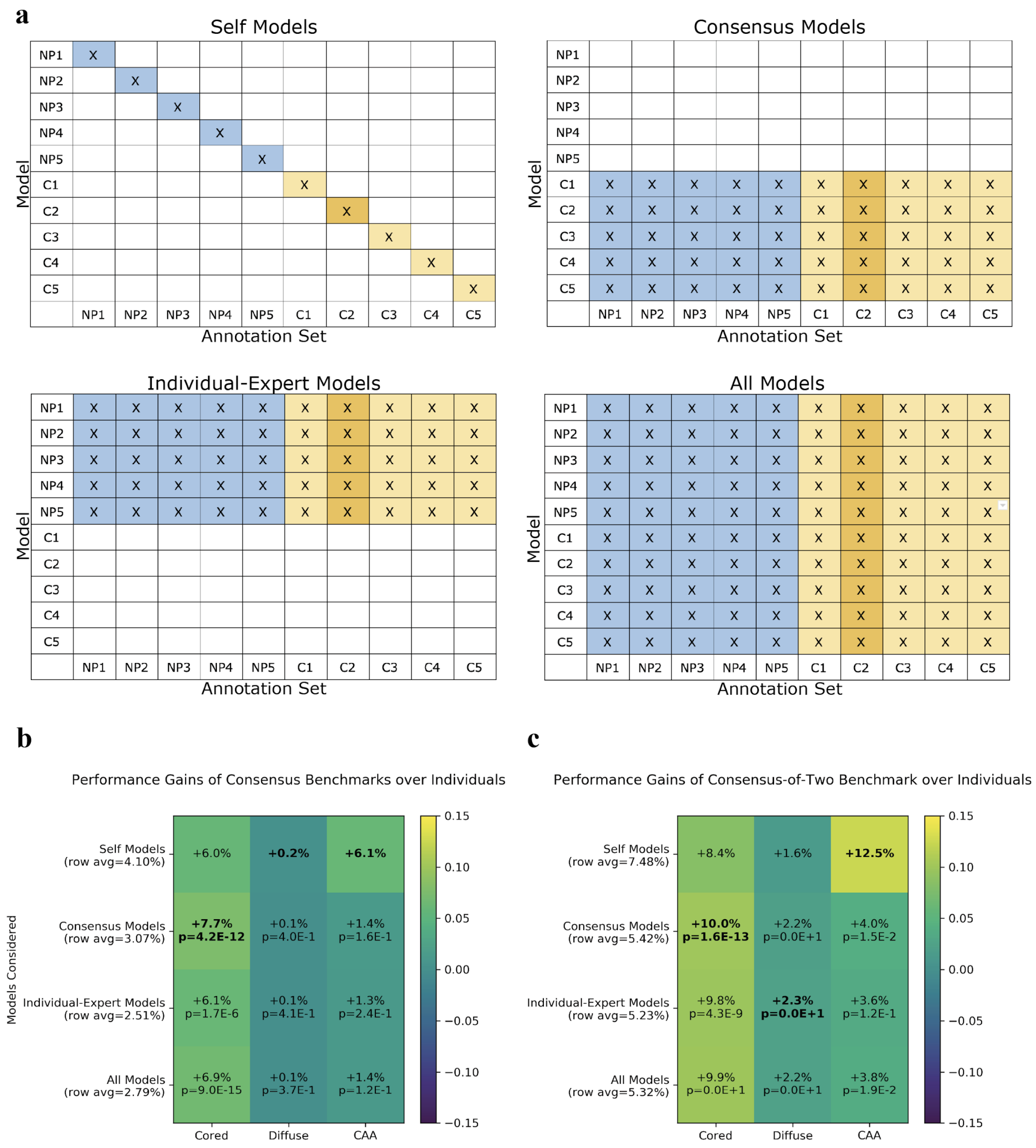


**Supplementary Figure 6: Consensus benchmarks facilitate higher model performance.** Corresponds to Figure 4, however this is comparing consensus versus individual-expert benchmarks as opposed to comparing models. (a) We compared the consensus benchmarks with the individual-expert benchmarks by evaluating subsets of models on every annotation benchmark. We compared the average performance of the models on the consensus-of-*n* benchmarks versus the average performance of the same models on the individual-expert benchmarks. We used four comparison schemes, selecting different models to evaluate ("self models," "consensus models," "individual-expert models," and "all models"). The first and most internally-consistent model scheme evaluated each individual-expert model according to the labels of its annotator (called "self models"). For consensus models, the annotation labels corresponded to labels derived from the matching consensus-of-*n* strategy. "Consensus models" evaluated every consensus model on every annotation set. "Individual-expert models" evaluated every individual-expert model on every annotation set. "All models" evaluated both consensus models and individual-expert models on every annotation set. The y-axis indicates the model, and the x-axis indicates the annotation set used to evaluate the model. The average AUPRC of the blue region (individual-expert benchmarks) is compared with the average AUPRC of the gold region (consensus benchmarks). The consensus-of-two is colored dark-gold for emphasis. (b) Depicts a heatmap of all consensus benchmarks vs individual benchmarks. We calculated p-values of the comparisons using a two-sample Z-test (Methods). P-values for the self-benchmark are not included because the sample size ($n$=20 comparisons) is not large enough to assign significance. The x-axis shows the Aβ class being evaluated, while the y-axis indicates the models that are evaluating the benchmark. We bolded the highest performance differential for each Aβ class. Regardless of what set of models we used, the annotations derived by a consensus-of-*n* strategy allowed for greater average model performance across all Aβ classes. (c) Depicts just the consensus-of-two benchmark versus the individual-expert benchmarks. For this consensus-of-two benchmark evaluation, only dark-gold regions in (a) corresponding to the consensus-of-two benchmark are compared to the blue region.

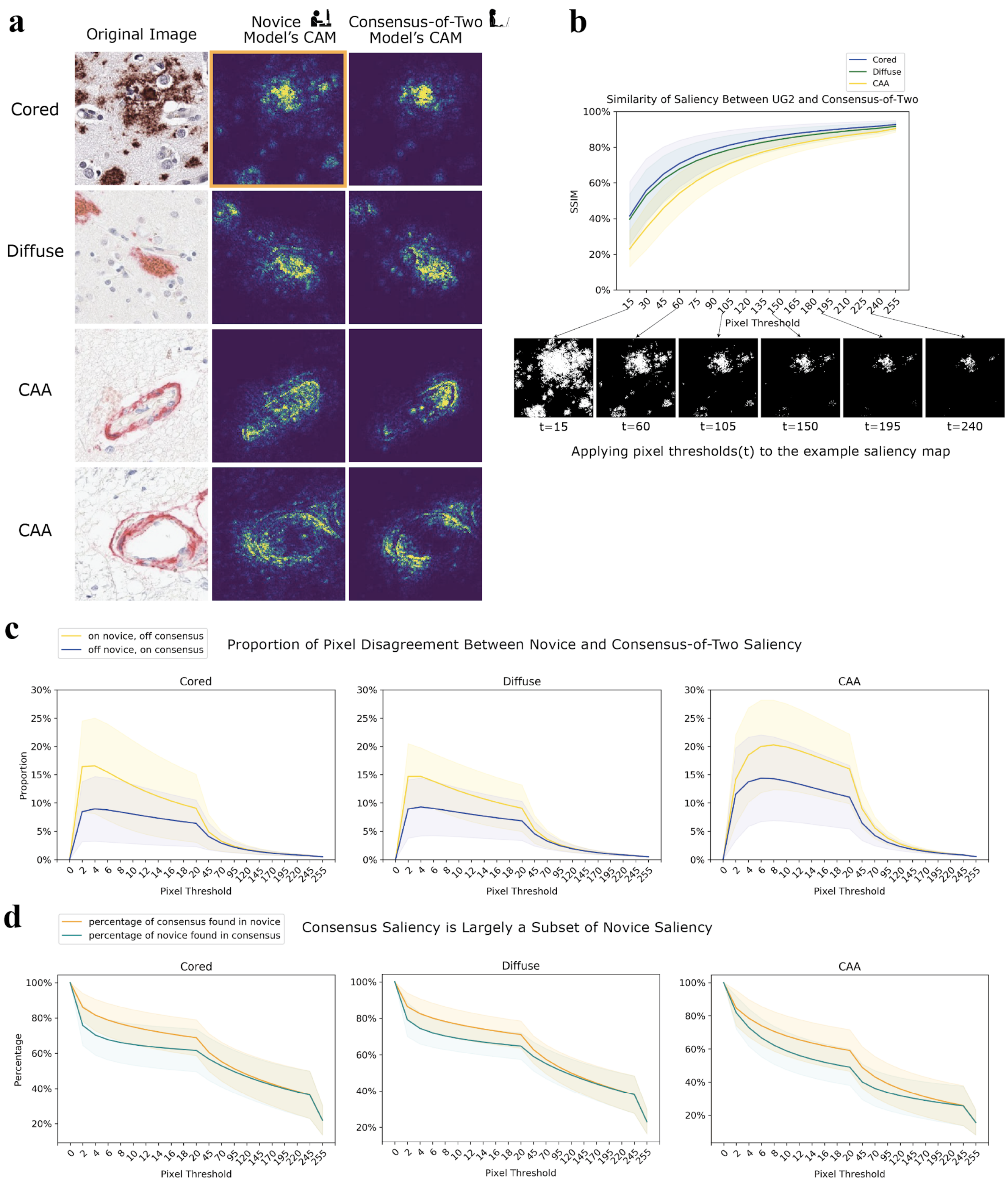


**Supplementary Figure 7: CAM analysis of UG2.** Corresponds to Figure 5, but for the second undergraduate novice. (a) Novice CAMs are more diffuse than expert CAMs. The original image (leftmost column), the CAM of the novice model trained on UG1's annotations (middle column), and the CAM of the consensus-of-two model (rightmost column). CAMs are plotted with a false-color map such that bright regions correspond to high intensity regions with high salience. (b) Although expert and novice CAMs differ, they converge on the same pixels. We progressively assess the structural similarity index (SSIM) between novice CAMs and consensus-of-two CAMs across the entire test set of images. The CAMs show the most similar salience by SSIM (y-axis) at the highest pixel thresholds as we increment the threshold (x-axis) used to binarize the images before comparison. Binarized examples are shown of one CAM from (a) (boxed in orange). (c) Comparing the novice CAMs and the consensus-of-two CAMs, we classify each pixel location into two categories: ON in the novice CAM and OFF in the corresponding consensus CAM (yellow), or OFF in the novice CAM and ON in the consensus CAM (blue). ON and OFF are determined by binarizing the images at pixel threshold *t* (x-axis). Y-axis shows the proportions at which these two cases occur. Zoomed inset highlights disagreement between CAMs. (d) Consensus CAM pixels are mostly contained within the novice CAM. The x-axis plots the varying pixel thresholds, while the y-axis plots the percent overlap of either how much of the consensus CAM pixels are a subset of the novice CAM (orange) or how much those of the novice CAM are a subset of the consensus (cyan).

**a**

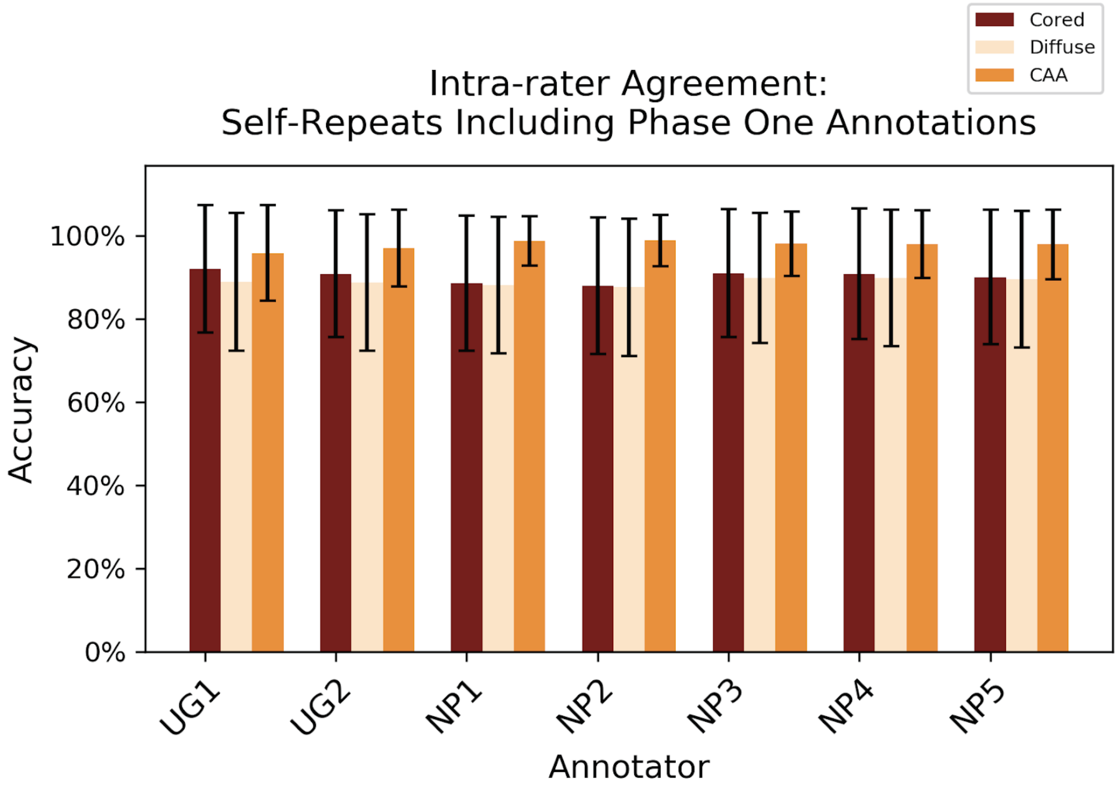

Intra-rater Agreement:
Self-Repeats Including Phase One Annotations
Cored
Diffuse
CAA
Accuracy
0%
20%
40%
60%
80%
100%
UG1
UG2
NP1
NP2
NP3
NP4
NP5
Annotator


**b**

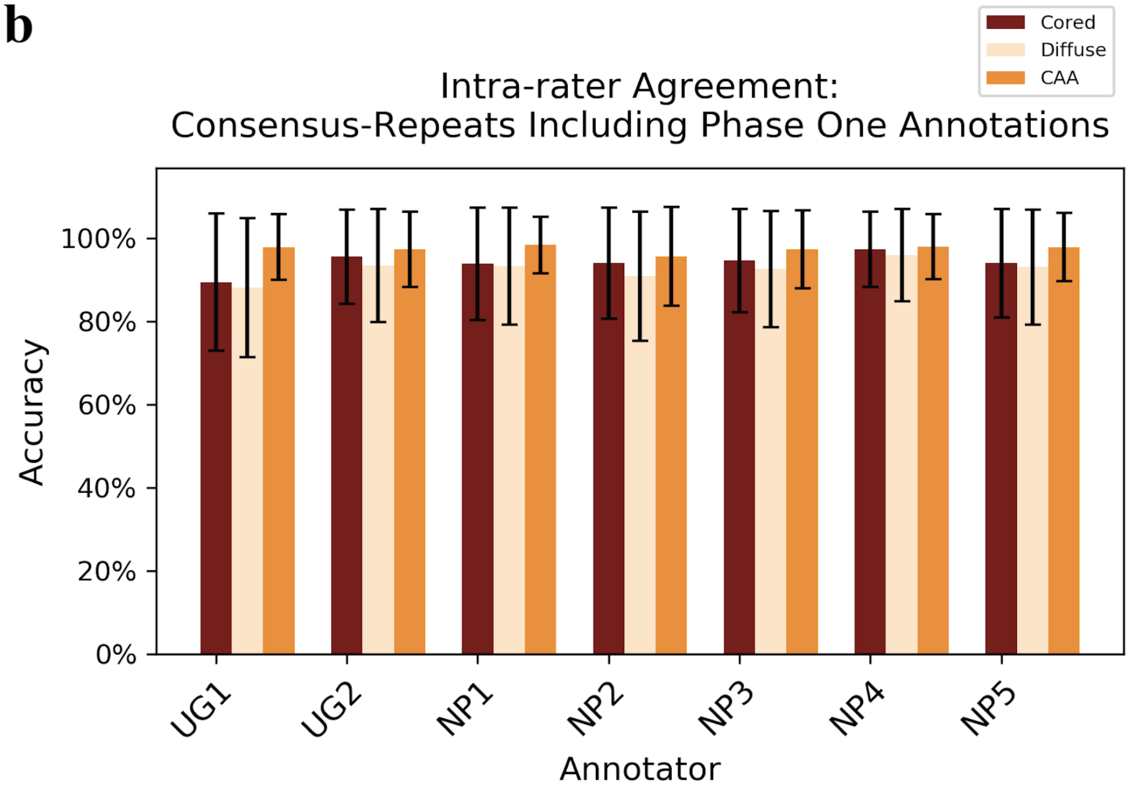

Intra-rater Agreement:
Consensus-Repeats Including Phase One Annotations
Cored
Diffuse
CAA
Accuracy
0%
20%
40%
60%
80%
100%
UG1
UG2
NP1
NP2
NP3
NP4
NP5
Annotator

**Supplementary Figure 8: Intra-rater agreement for self-enrichment and consensus-enrichment.** (a) Intra-rater agreement when only calculated on self-repeat images and phase one data. Novices achieved an average intra-rater agreement accuracy of 0.92 for cored, 0.89 for diffuse, and 0.97 for CAA. Experts achieved an average intra-rater agreement accuracy of 0.90 for cored, 0.89 for diffuse, and 0.98 for CAA. (b) Intra-rater agreement when only calculated on consensus-repeat images and phase one data. Novices achieved an average intra-rater agreement accuracy of 0.93 for cored, 0.91 for diffuse, and 0.98 for CAA. Experts achieved an average intra-rater agreement accuracy of 0.95 for cored, 0.94 for diffuse, and 0.97 for CAA.

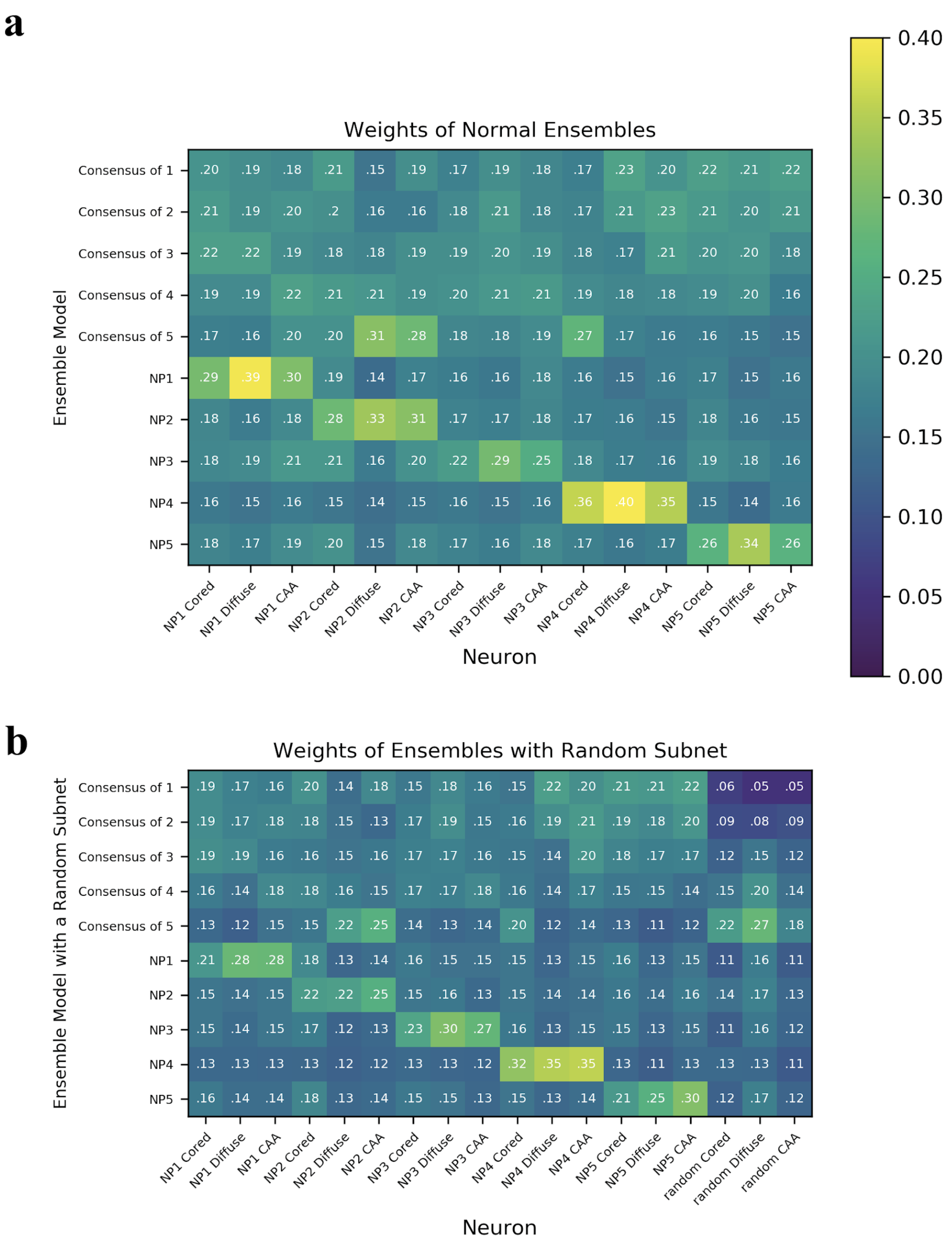


**Supplementary Figure 9: Ensemble weights.** The y-axis specifies the ensemble model, while the x-axis specifies the output neuron from the single CNN. Values are the softmaxed weights attached from each CNN's class output neurons to the final ensemble output layer. We apply a softmax function to the weights across each of the three amyloid classes. (a) depicts the normal ensembles (five single expert CNNs connected with an affine layer). (b) depicts the ensembles with a single random subnet (Five single expert CNNs and a random CNN connected with an affine layer). The “Random” neurons specify the class output neurons of the random CNN. The softmaxed weights provide a sense of relative importance of each CNN and how much effect each CNN has on the final output.

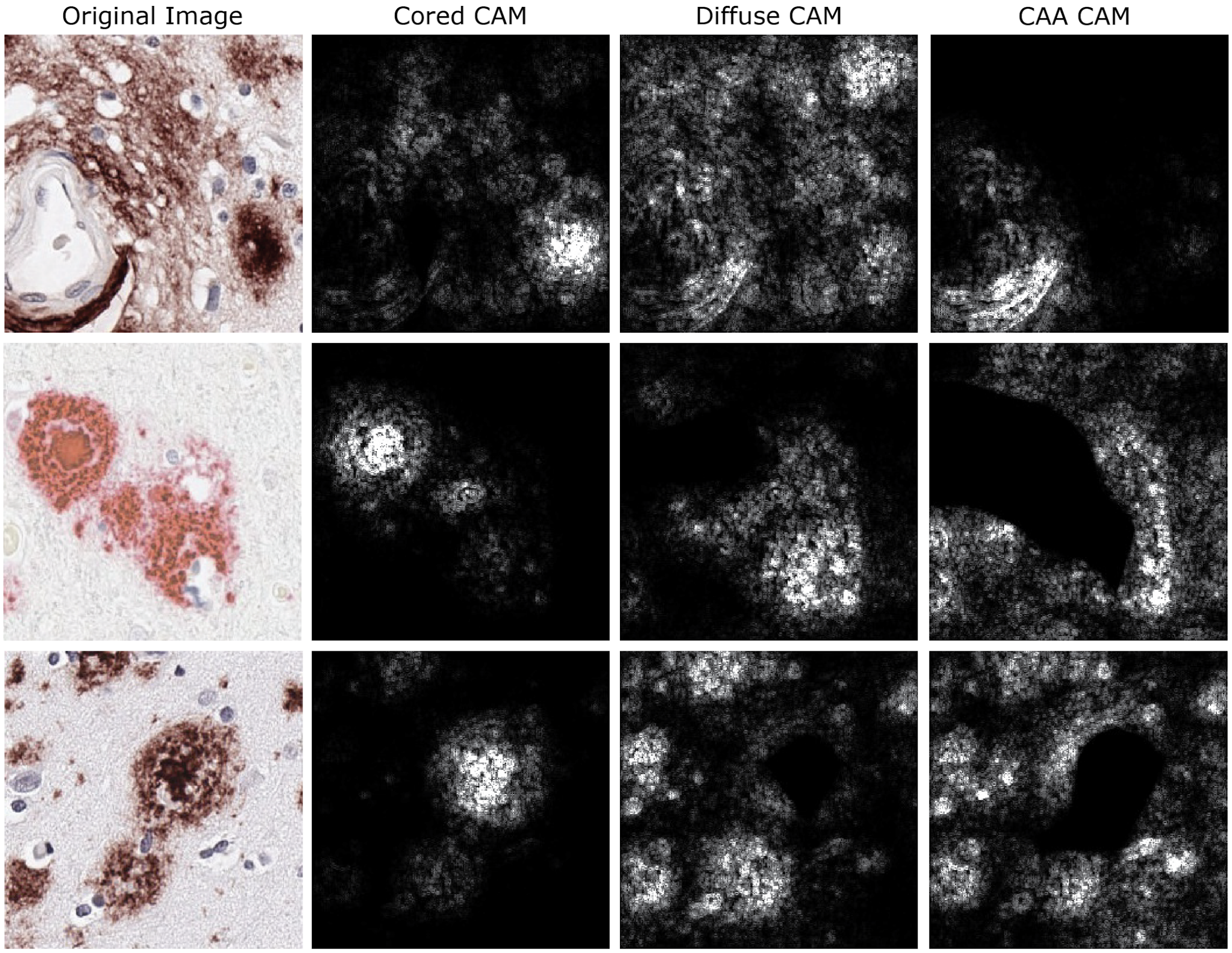


**Supplementary Figure 10: CAMs for each class**. We queried the consensus-of-two model to generate a CAM for each Aβ class (designated by column header). The leftmost column indicates the original image being analyzed. We generated a CAM for each class regardless of whether the image was positive for the class. Each image was labeled as positive by all five annotators (a full consensus-of-five)—the top row was positive for CAA, and the bottom two rows were positive for both cored and CAA.

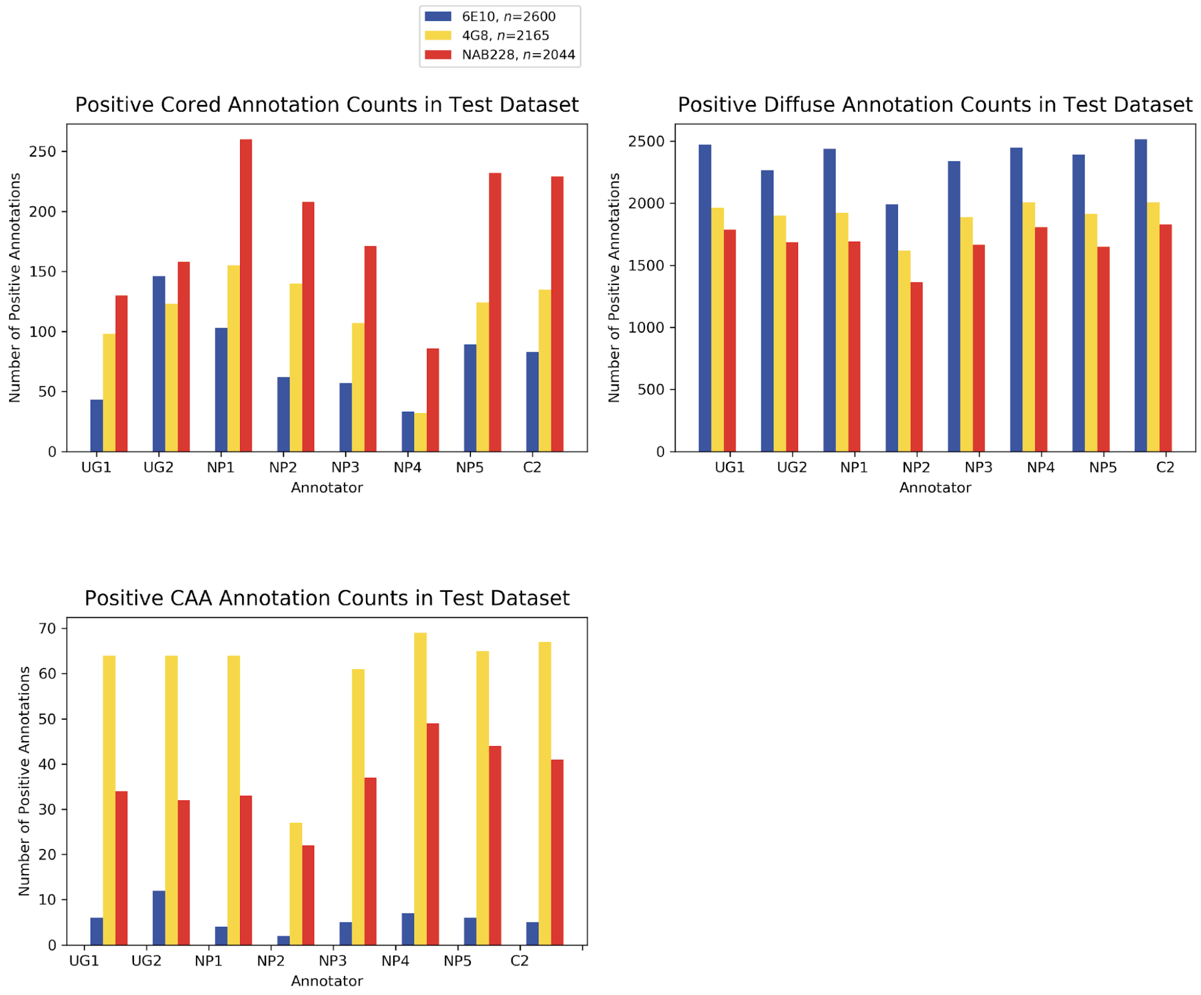


**Supplementary Figure 11: Distribution of positive annotations in the test set.** Each plot shows the number of positive annotations for a specific Aβ class in the test set. There are noticeable differences between stains, even within the same Aβ class. The CAA class with 6E10 is the most underrepresented in the test set.
